# Independent tuning of outer membrane fluidity and mechanics in Gram-negative bacteria

**DOI:** 10.64898/2026.09.17.752130

**Authors:** Jiawei Sun, Gvantsa Gutishvili, You He, Ryan Valdez, Handuo Shi, Petra Levin, Thomas J. Silhavy, James C. Gumbart, Steven Rutherford, Kerwyn Casey Huang

**Author notes:** These authors contributed equally (Gvantsa Gutishvili, You He).

## Abstract

The outer membrane (OM) of Gram-negative bacteria forms a protective barrier that combines selective permeability with mechanical load-bearing capacity, properties linked to its asymmetric bilayer structure with lipopolysaccharides (LPS) in the outer leaflet and phospholipids (PLs) in the inner leaflet. Unlike most PL bilayers, the OM typically exhibits limited lateral diffusion, resulting in a gel-like surface with spatially organized proteins and LPS. The molecular basis of this physical state and its relationship with envelope mechanics remain unclear. Here, we show that increasing PL levels in the outer leaflet or truncating LPS core oligosaccharides increases OM fluidity by disrupting LPS packing. In contrast, reduced LPS abundance or disruption of divalent cation-mediated crosslinking primarily reduces OM stiffness with little effect on fluidity. Thus, OM fluidity and mechanical stiffness can be tuned independently through distinct molecular interactions. This separation of physical control mechanisms provides a framework for understanding how Gram-negative bacteria modulate OM properties during environmental adaptation and envelope homeostasis.

## Introduction

The outer membrane (OM) of Gram-negative bacteria provides essential protection for growth and survival^1^. The OM is an asymmetric bilayer composed of lipopolysaccharides (LPS) in the outer leaflet, phospholipids (PLs) in the inner leaflet, and transmembrane β-barrel proteins that occupy a substantial fraction of the cell surface^2^. This architecture supports multiple functions, including acting as a permeability barrier and a mechanical load-bearing structure^3^. LPS in the outer leaflet restricts entry of potentially harmful molecules, including many antibiotics^1,4,5^.

Complementing this chemical barrier, recent studies showed that the OM exhibits stiffness comparable to the rigid peptidoglycan (PG) cell wall, enabling it to resist mechanical stresses arising from changes in external osmolarity^6^. The OM is mechanically coupled to the cell wall by abundant linker molecules^7–12^, allowing the periplasm to function as an integrated mechanical unit that buffers turgor pressure during hypoosmotic shocks and tolerates osmolyte efflux from the cytoplasm^13^. During cell growth, the biogenesis of PG and the OM must therefore be coordinated to ensure synchronous expansion^14,15^. The OM also contributes to maintaining cell shape, as mutants with reduced OM protein levels exhibit morphological defects^16^, whereas strengthening the OM can restore rod shape in certain mutants with impaired cell wall synthesis^17^.

The physical properties of the OM are shaped by interactions among LPS, proteins, and PLs^3,18–20^. The OM exhibits pronounced spatial heterogeneity^21–23^ driven by preferential insertion of OM material at mid-cell^24^. This biased insertion is facilitated by inhibition of Bam complexes, which are responsible for OM protein insertion, by tetrapeptides present in mature PG near the cell poles^15^. As a result, newly synthesized OM components accumulate near the midcell while older proteins are displaced toward the poles^24^, generating spatial organization across the cell surface. Such patterning may promote turnover of OM proteins during growth^24^ and produce phenotypic heterogeneity between daughter cells^25^. At smaller scales, OM proteins form clusters visible via fluorescence microscopy^21,24,26^, while LPS-rich domains and lattice-like porin networks have been observed using atomic force microscopy^2^. Maintaining this spatial organization requires that surface molecules diffuse slowly enough to prevent homogenization by passive diffusion. Indeed, both OM proteins and LPS exhibit limited lateral mobility^21,27–31^, in contrast to the fluid-like behavior of most lipid bilayers, including the bacterial inner membrane (IM)^32,33^. Increased OM fluidity has been associated with compromised membrane integrity and increased permeability^30,34–36^, suggesting that permeability, fluidity, and mechanical strength may arise from common molecular interactions. However, despite the physiological importance of OM fluidity and spatial organization, the molecular mechanisms that determine OM fluidity, particularly how they relate to OM mechanics, remain poorly understood.

Here, we quantify LPS diffusion using fluorescence recovery after photobleaching (FRAP) of fluorescently labeled LPS (**Methods**)^30,37^. Because LPS constitutes a dominant component of the outer leaflet and coordinates the molecular interaction network at the OM surface, the lateral diffusion of labeled LPS provides a useful readout of the effective fluidity of the OM. Measuring LPS diffusion in *Escherichia coli* across a broad set of chemical and genetic perturbations revealed two distinct mechanisms controlling OM fluidity. Mutants with reduced levels of OM proteins exhibited large, growth-phase-dependent increases in fluidity associated with PL accumulation in the outer leaflet. In low-salt conditions, truncation of LPS core oligosaccharides also increased OM fluidity, likely by weakening intermolecular interactions between neighboring LPS molecules and disrupting their ordered packing. Strikingly, these perturbations altered OM fluidity without proportionally affecting OM stiffness. By contrast, reducing LPS abundance or acutely removing divalent cations that mediate LPS crosslinking strongly decreased OM stiffness while causing only modest changes in fluidity. Together, these results demonstrate that OM fluidity and mechanical stiffness can be tuned independently through distinct molecular interactions.

## Results

### PL accumulation increases LPS mobility

As an asymmetric bilayer rich in transmembrane porins and LPS, the OM exhibits physical properties distinct from those of the IM, likely arising from strong interactions among LPS and abundant OM proteins^3^. Previous studies showed that LPS crosslinking by divalent cations^6^ and the presence of OM proteins^16^ contribute to the load-bearing capacity of the *E. coli* OM. We therefore tested whether interactions among LPS and OM proteins determine OM fluidity, with stronger molecular interactions expected to reduce molecular mobility.

To quantify LPS diffusion, we measured fluorescence recovery using FRAP following metabolic labeling of LPS through an azide-alkyne reaction targeting 3-deoxy-D-manno-oct-2-ulosonic acid (Kdo) with Rhodamine 110 (**Fig. 1a; Methods**)^30,37^. Fluorescence recovery profiles of individual cells were compared with simulated one-dimensional diffusion profiles based on the initial photobleaching pattern to estimate diffusion coefficients (**Fig. 1a; Methods**).

**Figure 1:**
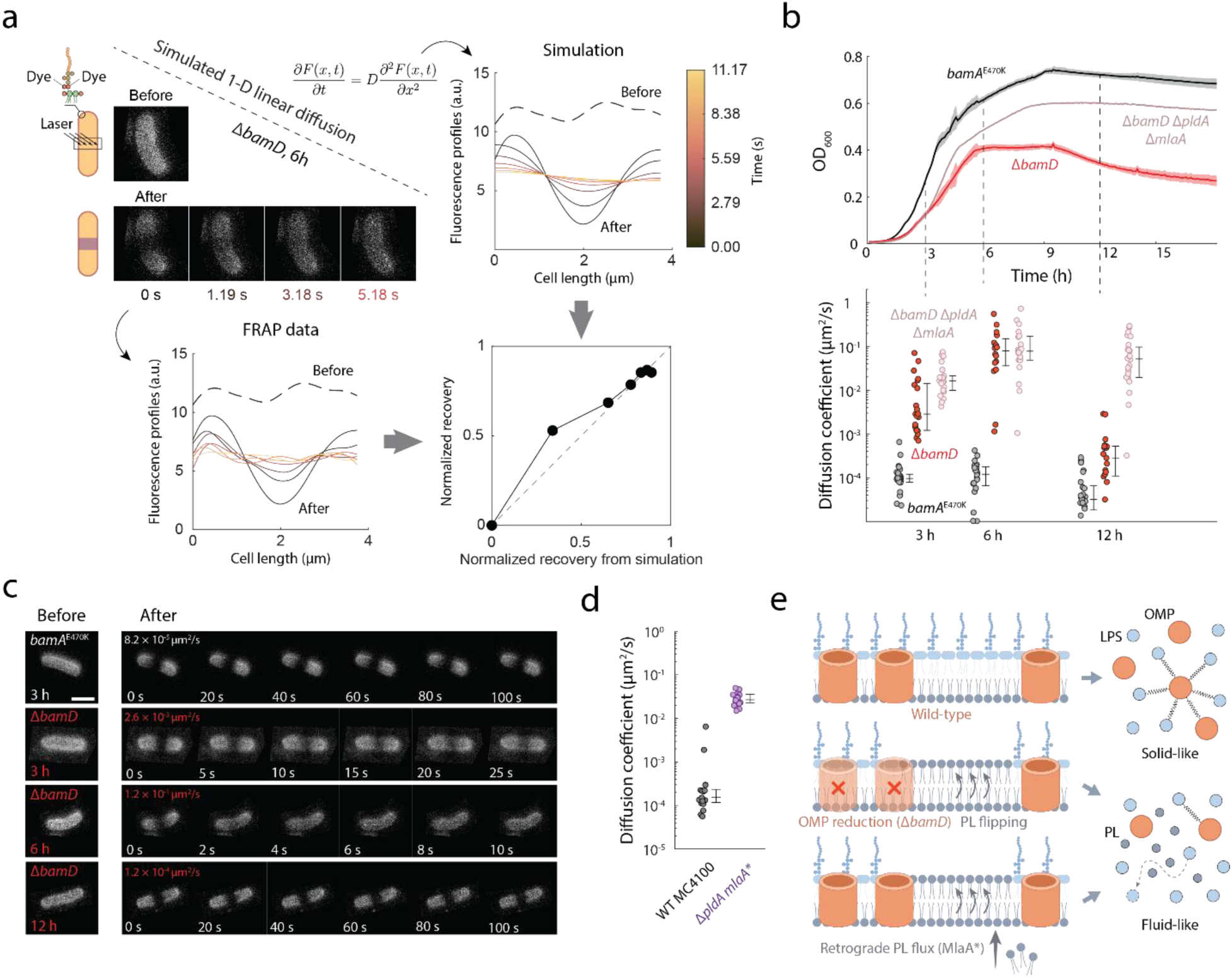
Insertion of PLs into the outer leaflet fluidizes the OM by disrupting LPS packing. a) Schematic of the FRAP assay used to measure LPS lateral diffusion. LPS molecules were labeled via Kdo-N_3_ incorporation and azide-alkyne click chemistry. FRAP was quantified by comparing experimental fluorescence recovery profiles to simulated one-dimensional diffusion profiles based on the diffusion equation (**Methods**). b) Δ*bamD* and Δ*bamD* Δ*pldA* Δ*mlaA* cells, which have strongly reduced levels of OM proteins^16^, exhibit increased OM fluidity during log-phase and early stationary-phase growth in LB medium, as measured by the estimated LPS diffusion coefficient (**Methods**), coincident with PL accumulation in the OM outer leaflet. In late stationary phase (12 h), when PLs are removed from the outer leaflet, fluidity in Δ*bamD* cells returns to levels similar to the wild-type-like parent strain *bamA^E470K^* (gray), whereas fluidity remains elevated in Δ*bamD* Δ*pldA* Δ*mlaA*, which retain high PL levels. Upper: growth curves are shown as mean ± 1 standard deviation (s.d.) from three technical replicates. Lower: distributions of diffusion coefficients. Center line indicates medians, and error bars indicate the 25^th^ and 75^th^ percentiles. *N* > 15 cells were tested per condition. c) Representative time-lapse images showing fluorescence recovery after photobleaching in *bamA^E470K^*, Δ*bamD*, and Δ*bamD* Δ*pldA* Δ*mlaA* cells at different growth phases in LB medium. Scale bar: 2 µm. d) OM fluidity is increased in log-phase *mlaA*\* Δ*pldA* cells grown in LB, which accumulate PLs in the outer leaflet without direct disruption of OM protein assembly. Center lines indicate medians, and error bars indicate the 25^th^ and 75^th^ percentiles. *N* > 17 cells were tested per condition. e) Model illustrating OM fluidization caused by PL accumulation in the outer leaflet. Increased PL content, resulting from reduced OM protein levels or enhanced PL flux, disrupts the tight packing of LPS molecules.

To investigate the contribution of OM proteins in OM fluidity, we examined a previously characterized Δ*bamD* mutant. Deletion of *bamD* causes a global reduction in OM proteins, with PLs filling the resulting void by flipping from the inner leaflet to the outer leaflet during log-phase growth^16^. In stationary phase, the Δ*bamD* OM becomes mechanically destabilized due to the activities of OM phospholipase A and the Mla system^38–43^, which remove PLs from the outer leaflet and eventually cause OM rupture in a fraction of cells^16^. Thus, outer-leaflet PL levels are expected to increase during growth and decrease in stationary phase. Consistent with this model, Δ*bamD* cells (**Table S1**) exhibited a ∼1000-fold increase in LPS diffusion after 6 h of growth in LB medium (early stationary phase) compared with cells immediately back-diluted from a 12-h culture representing late stationary phase and the onset of regrowth (**Fig. 1b,c**). Fluorescence recovery kinetics closely resembled normal diffusion (**Fig. S1**). In contrast, the *bamA^E470K^* parent strain exhibited similarly low LPS diffusion across growth phases, comparable to late stationary-phase Δ*bamD* cells (**Fig. 1b,c**).

These growth phase-dependent changes in OM fluidity in Δ*bamD* cells mirrored previously characterized dynamics of PL accumulation in the outer leaflet during log phase and subsequent PL removal during stationary phase^16^, suggesting that outer-leaflet PL abundance is an important determinant of OM fluidity. To test this hypothesis, we quantified LPS diffusion in a Δ*bamD* Δ*pldA* Δ*mlaA* mutant (**Table S1**), in which loss of PldA-mediated PL degradation and Mla-dependent PL removal promotes retention of PLs in the OM outer leaflet^16^. In this triple mutant, OM fluidity remained high across all growth phases (**Fig. 1b**). Further supporting this interpretation, a Δ*bamB* mutant, which has a milder defect in OM protein folding than Δ*bamD*^44,45^, exhibited intermediate, growth phase-dependent OM fluidization (**Fig. S2**), consistent with lower levels of PL accumulation in the outer leaflet.

Because Δ*bamD* simultaneously reduces OM protein abundance and increases outer-leaflet PLs, we next asked whether PL accumulation could increase OM fluidity without direct disruption to OM protein assembly. Unlike *bam* mutants, which disrupt the BAM complex and globally reduce OMP levels, the gain-of-function allele *mlaA*\* primarily perturbs lipid asymmetry by promoting PL accumulation in the outer leaflet^46^. In an *mlaA*\* Δ*pldA* mutant, in which PLs are driven into and retained in the outer leaflet, OM fluidity was high (**Fig. 1d**). This is consistent with the disruption of LPS-OM protein packing in Δ*mlaA* Δ*pldA* cells previously observed using atomic force microscopy^2^. These results demonstrate that outer-leaflet PL accumulation can drive increased LPS diffusion even when the primary perturbation does not directly disrupt OM protein assembly, supporting PL-mediated disruption of LPS packing as a major determinant of OM fluidization.

### LPS core oligosaccharides restrict OM fluidity under low-salt conditions

LPS is an amphipathic molecule composed of a lipid moiety (lipid A), core oligosaccharides, and, in some strains, a long polysaccharide chain termed the O-antigen^47^ (in most *E. coli* laboratory strains, including the K-12-derived strains used in this study, the O-antigen is absent). LPS contains multiple negatively charged phosphate groups that promote formation of an ordered LPS network through salt bridges mediated by divalent cations^20^. Core oligosaccharides further mediate interactions between LPS molecules and contribute to OM mechanical stiffness^48^.

Based on this structure, we hypothesized that shortening the core oligosaccharides would weaken interactions between neighboring LPS molecules and thereby increase OM fluidity relative to wild-type cells. To test this hypothesis, we measured LPS diffusion in mutants defective in core oligosaccharide synthesis (**Fig. 2a,b and S3**). The *waa* genes encode enzymes that sequentially catalyze addition of sugars to the LPS core^49^, and deletion of specific *waa* genes produces LPS molecules truncated to varying extents.

**Figure 2:**
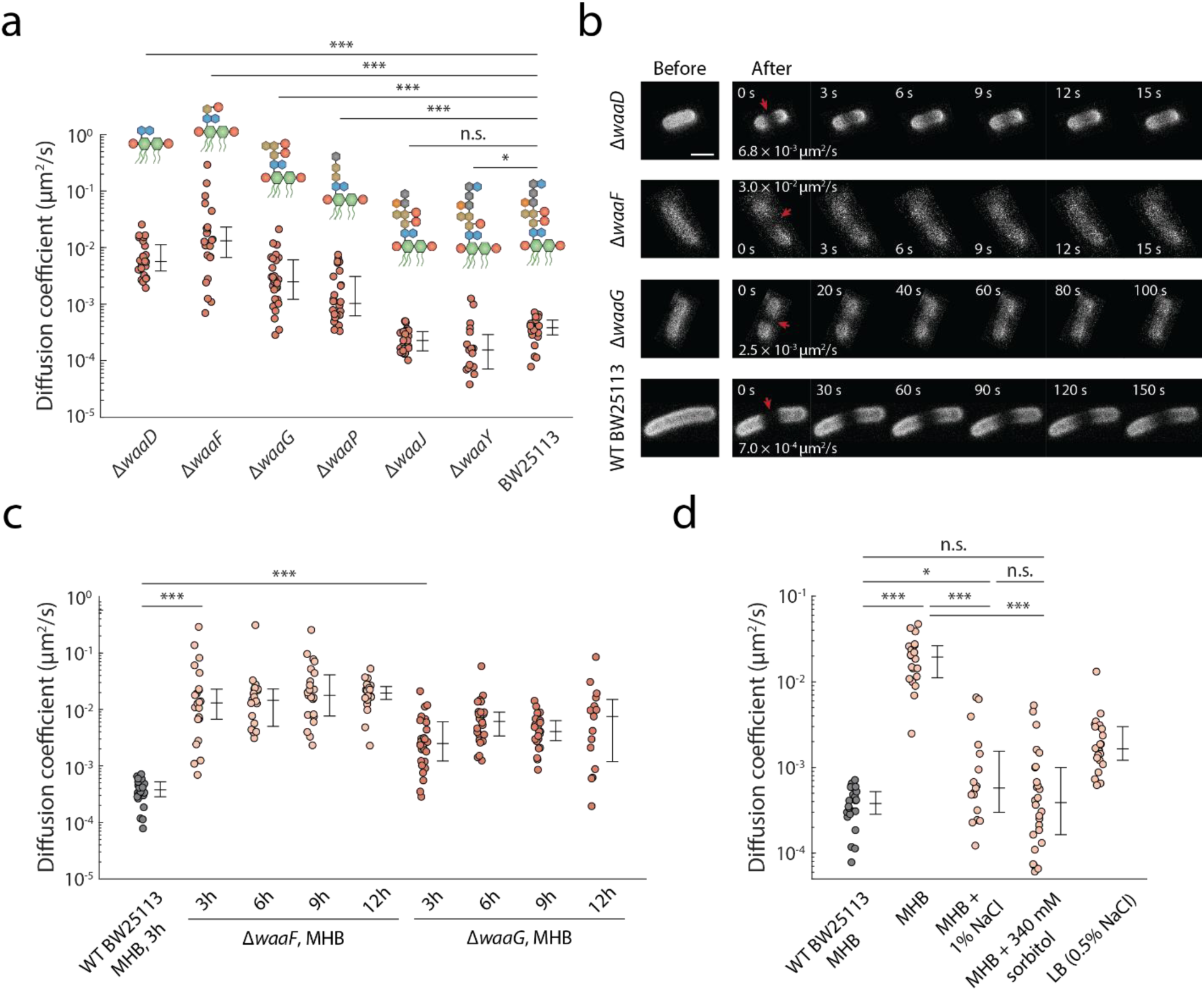
OM fluidity depends on the length of LPS core oligosaccharides. a) Genetic truncation of LPS core oligosaccharides progressively increased OM fluidity in NaCl-deficient MHB medium, with the magnitude of the effect depending on the remaining oligosaccharide length. Cells were grown for 3 h prior to measurement. Center lines indicate medians, and error bars indicate the 25^th^ and 75^th^ percentiles. *N* > 18 cells were tested per condition. *p* values were calculated using Welch’s t-test on log_10_-transformed diffusion coefficients. ***: *p* < 0.001; *: *p* < 0.05; n.s.: *p* > 0.05. b) Representative time-lapse images showing fluorescence recovery after photobleaching in wild-type, Δ*waaF*, Δ*waaD*, and Δ*waaG* cells grown in MHB. c) OM fluidity was largely independent of growth phase in Δ*waaF* and Δ*waaG* cells, with no statistically significant differences detected across time points. Center lines indicate medians, and error bars indicate the 25^th^ and 75^th^ percentiles. *N* > 14 cells were tested per condition. *p* values were calculated using Welch’s t-test on log_10_-transformed diffusion coefficients. ***: *p* < 0.001; n.s.: *p* > 0.05. d) Supplementation of MHB with 1% NaCl or 340 mM sorbitol (osmotically equivalent to 1% NaCl) restored low OM fluidity in Δ*waaD* cells. Growth in LB (Lennox; 0.5% NaCl) partially restored low fluidity. Diffusion coefficients are shown as box plots; center lines indicate medians, and error bars indicate the 25^th^ and 75^th^ percentiles. *N* > 16 cells per condition. *p* values were calculated using Welch’s t-test on log_10_-transformed diffusion coefficients. ***: *p* < 0.001; *: *p* < 0.05; n.s.: *p* > 0.05.

A previous study reported that Δ*waaD* exhibits increased diffusion of fluorescent pyrenedecanoic acid probes incorporated into the OM when cells are grown in the low-salt rich medium MHB, compared to MHB supplemented with 1% NaCl^50^. Motivated by this observation, we quantified LPS diffusion in *waa* mutants^51^ (**Table S1**) grown in MHB. Consistent with a model in which reduced LPS-LPS interactions increase OM fluidity, the extent of fluidization depended on the remaining length of the core oligosaccharides (**Fig. 2a,b**). Δ*waaD* and Δ*waaF* mutants, which possess the shortest inner cores^52–54^, exhibited the highest fluidity (**Fig. 2a**), whereas Δ*waaJ*, whose LPS structure is only modestly truncated, showed no significant increase (**Fig. 2a**). These results suggest that longer core oligosaccharides restrict LPS mobility, potentially by strengthening interactions between neighboring LPS molecules.

Δ*waaP* cells also exhibited increased fluidity, which may reflect loss of phosphate groups and/or impaired synthesis of the outer core^55^. Unlike the *bam* mutants, increased OM fluidity in Δ*waaF* and Δ*waaG* cells was independent of growth phase (**Fig. 2c**), consistent with weakened interactions between truncated core oligosaccharides rather than growth phase-dependent PL insertion into the outer leaflet.

Environmental salt conditions strongly influenced these effects. In LB (Lennox) or MHB supplemented with 1% NaCl, Δ*waaD* cells exhibited substantially lower fluidity than in unsupplemented MHB (**Fig. 2d**), consistent with previous observations of salt dependence in wild-type cells^50^. This reduction could arise either from ionic effects that strengthen LPS-LPS interactions or from increased environmental osmolarity. To distinguish between these possibilities, we measured LPS diffusion in NaCl-deficient MHB supplemented with 340 mM sorbitol, which matches the osmotic strength of MHB+1% NaCl without introducing additional ions. Sorbitol restored low fluidity (**Fig. 2d**), indicating that the effects of NaCl arise primarily from increased external osmolarity rather than ionic interactions alone. These results suggest that higher osmolarity restricts LPS diffusion, consistent with increased packing or confinement of OM components.

To understand how truncation of LPS core oligosaccharides affects LPS motility and spatial packing, we performed all-atom molecular dynamics (MD) simulations of OM patches containing 34 LPS molecules derived from either wild-type or Δ*waaF* cells (**Fig. S4a; Methods**). These simulations revealed that LPS oligosaccharides interact primarily through direct hydrogen bonds and crosslinking mediated by divalent cations (**Fig. 3a,c**). Truncation of the core oligosaccharides reduced these interactions (**Fig. 3b,d**), relieving local confinement of LPS molecules and increasing local mobility (**Fig. 3e,f**). Although the simulation timescale (5 µs) was much shorter than the estimated time (∼1 ms) required for an LPS molecule with diffusion constant *D*∼10^-3^ µm^2^/s to traverse a distance comparable to its lateral size (∼1.5 nm), diffusion coefficients estimated from mean-squared displacement were consistent with experimentally measured values (**Fig. S4b,c; Methods**).

**Figure 3:**
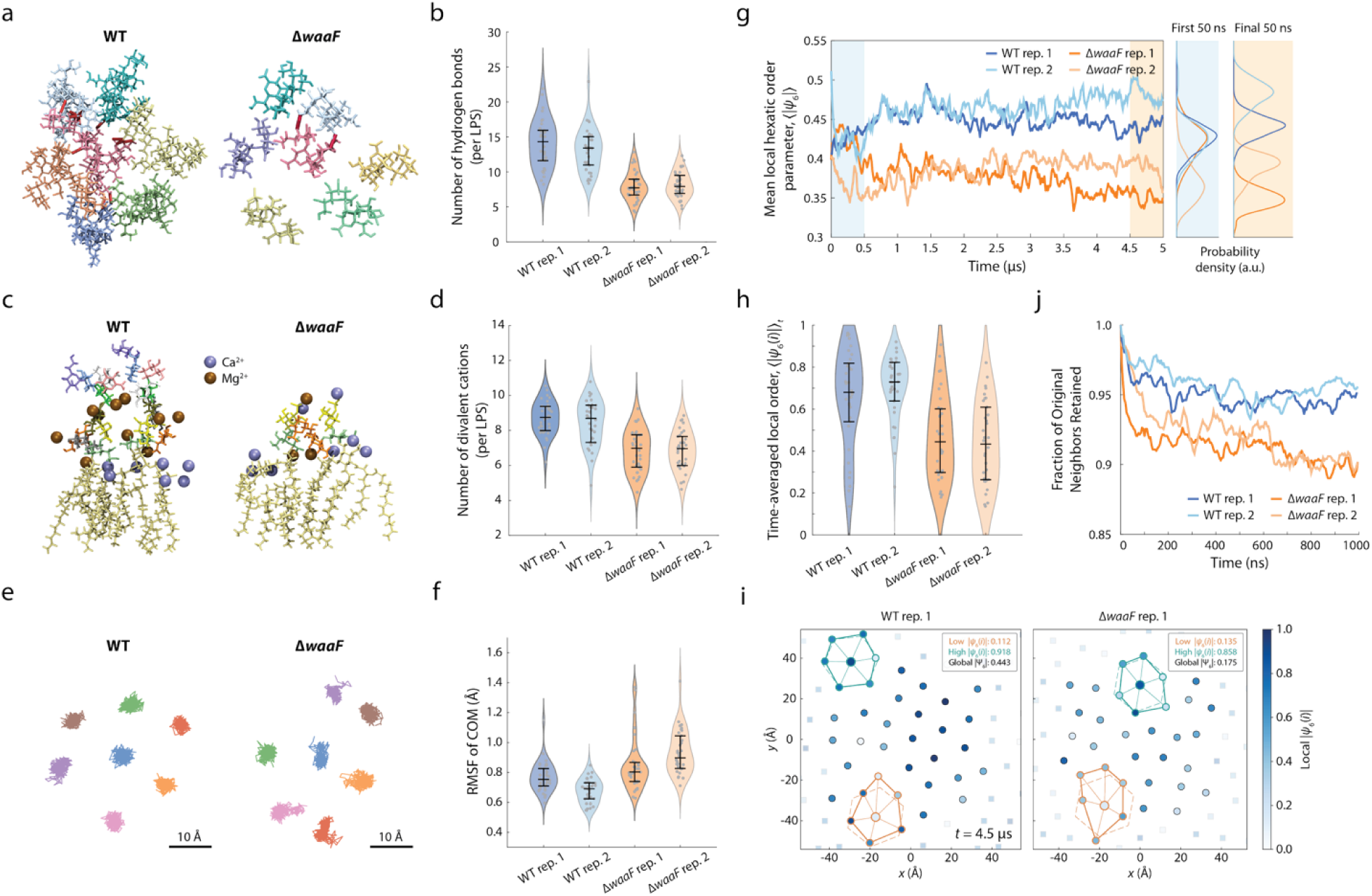
Truncation of LPS core oligosaccharides weakens intermolecular interactions, disrupts packing, and increases local mobility in MD simulations. a) Representative LPS molecule and hydrogen bonds with its six nearest neighbors in wild-type and Δ*waaF* systems. Truncated Δ*waaF* LPS forms fewer hydrogen bonds with neighboring LPS molecules. Each LPS is shown in a different color, and thick red lines denote hydrogen bonds. b) Truncation reduces the number of hydrogen bonds formed by individual LPS molecules with neighboring LPS, quantified over the final 1 µs of simulation. Each violin plot represents an independent simulation. Data points correspond to individual LPS molecules; center lines indicate medians, and error bars indicate the 25^th^ and 75^th^ percentiles. c) Representative LPS molecules and associated divalent cations in wild-type and Δ*waaF* systems. Mg^²2^ and Ca^²2^ ions within 4 Å of LPS are shown, illustrating reduced association of divalent cations with truncated LPS. d) Truncation reduces the number of divalent cations associated with individual LPS molecules over the final 1 µs of simulation, illustrating the weakening of interactions that stabilize neighboring LPS molecules. Violin plots show distributions for individual LPS molecules in each independent simulation; center lines indicate medians, and error bars indicate the 25^th^ and 75^th^ percentiles. e) Two-dimensional center-of-mass (COM) trajectories during the final 1 µs of simulation of lipid A for a representative LPS molecule and its six nearest neighbors, as determined from the initial configuration. The greater spatial excursions in Δ*waaF* illustrate increased local motion following core truncation. f) Truncated Δ*waaF* LPS exhibits increased local lateral motion, reflected by higher root-mean-square fluctuation (RMSF) of lipid A COM positions over the final 1 µs of simulation. Data points correspond to individual LPS molecules; center lines indicate medians, and error bars indicate the 25^th^ and 75^th^ percentiles. Two independent simulations were analyzed for each genotype. g) LPS core truncation reduces local packing order. Time courses of the mean local hexatic order parameter, 〈|*ψ*_6_|〉, which reports the angular organization of nearest neighbors around each LPS molecule (**Methods**), show that wild-type and Δ*waaF* LPS relax into distinct packing states, with Δ*waaF* maintaining lower local order. Shaded regions indicate the initial and final 50-ns intervals used to illustrate the distributions at right. h) Time-averaged local hexatic order for individual LPS molecules over the final 1 µs of simulation, 〈|*ψ*_6_(*i*)|〉*_t_*, confirms reduced local packing order in Δ*waaF* relative to wild type. Data points correspond to individual LPS molecules; center lines indicate medians, and error bars indicate the 25^th^ and 75^th^ percentiles. i) Representative endpoint configurations of wild-type and Δ*waaF* LPS layers colored by local |*ψ*_6_(*i*)|, illustrating the greater local packing order of wild-type LPS. Circles indicate molecules within the primary simulation box and squares indicate periodic images. Insets show representative central LPS molecules and their nearest-neighbor networks, with solid and dashed lines representing the configuration at *t*=4.5 µs and a perfect hexagonal lattice, respectively. j) Core truncation accelerates rearrangement of local LPS neighborhoods. The fraction of the original six nearest neighbors retained around each LPS molecule decreases more rapidly in Δ*waaF* than in wild-type, indicating that intact core oligosaccharides stabilize local LPS coordination shells and restrict neighbor exchange.

We next tested whether weakened interactions disrupt the organization of the LPS layer. In wild-type simulations, LPS molecules formed locally ordered hexagonal arrays, whereas Δ*waaF* simulations showed more disordered configurations across independent simulations (**Fig. 3g-i**), quantified by a lower local hexatic order parameter measuring nearest-neighbor angular alignment (**Methods**). Packing disruption also propagated to larger scales: Δ*waaF* LPS exhibited reduced long-range order, reflected in a lower global hexatic order parameter (**Fig. S4d; Methods**). To assess whether structural disorder was associated with a more dynamic local environment, we tracked neighbor exchange over time (**Methods**). Intermolecular contacts persisted significantly longer in wild-type simulations (**Fig. 3j**), demonstrating that core truncation destabilizes the local LPS lattice and accelerates molecular rearrangement. Together, these results indicate that intact core oligosaccharides stabilize local LPS organization by enhancing packing order and suppressing rapid molecular rearrangement, likely within a broader OMP-LPS interaction network.

### OM fluidity is largely insensitive to LPS abundance or divalent-cation crosslinking

Because LPS is a prominent component of the OM outer leaflet, reductions in LPS abundance might be expected to weaken intermolecular interactions and thereby increase OM fluidity. To test this possibility, we examined mutants with impaired LPS transport. LPS is synthesized in the cytoplasm and transported to the OM via the Lpt system^56^. The *imp4213* allele of *lptD* reduces LPS transport by impairing the function of LptD^44,45^, which together with LptE inserts LPS into the OM. Although *imp4213* (**Table S1**) reduces OM stiffness^6^ and increases permeability^44,45^, we observed only minimal increases in LPS diffusion in LB (**Fig. 4a**) and modest (∼10-fold) increases in MHB (**Fig. S5**). In *imp4213* cells, decreased LPS transport renders the OM permeable to hydrophobic compounds^44^ due to the formation of PL-rich domains that likely explain the modest increase in fluidity. Similarly, CRISPRi inhibition of LPS transport (*lptC*) or synthesis (*lpxC*; **Table S1**) resulted in only slight increases in fluidity (**Fig. 4a**) despite large reductions in LPS levels, in contrast to the substantial fluidization caused by *bamD* deletion (**Fig. 1b**).

**Figure 4:**
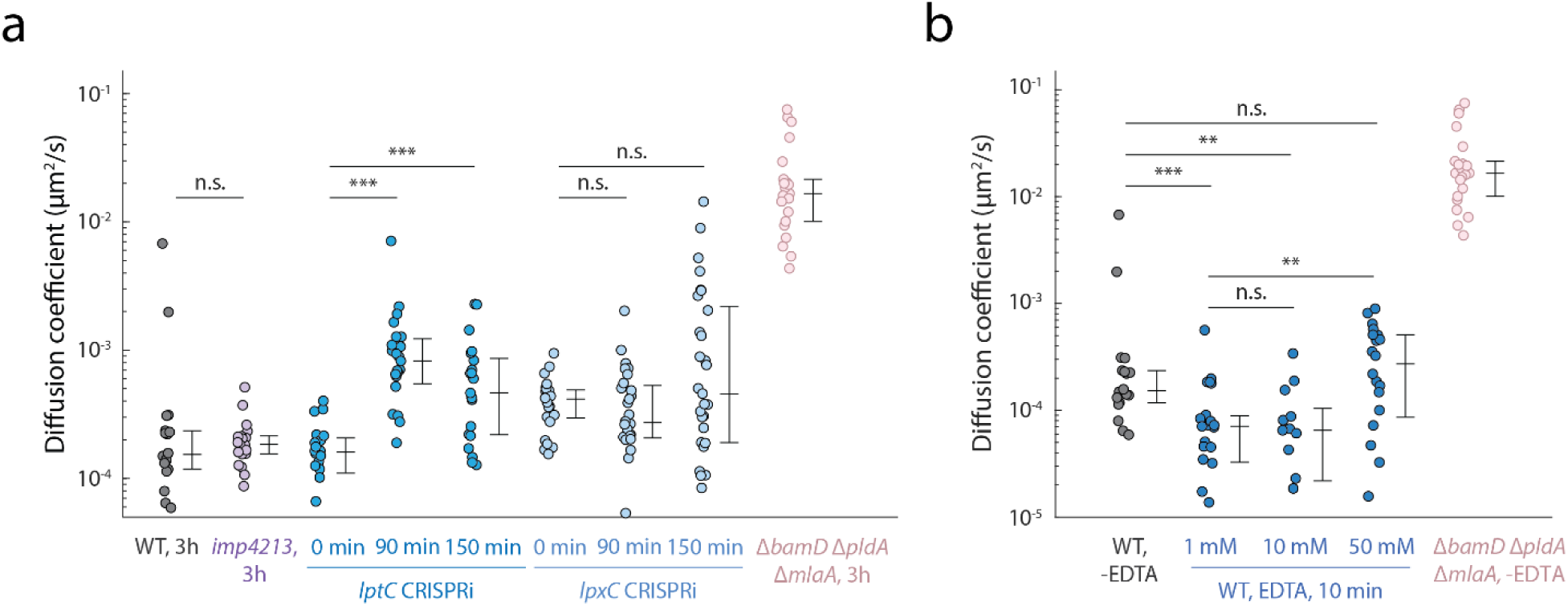
Disrupting LPS crosslinking or reducing LPS levels causes only modest increases in OM fluidity. a) Reducing LPS levels through the *imp4213* allele or CRISPRi inhibition of *lpxC* or *lptC* results in only modest increases in OM fluidity compared with PL accumulation in the OM outer leaflet (Δ*bamD* Δ*pldA* Δ*mlaA* cells). Center lines indicate medians, and error bars indicate the 25^th^ and 75^th^ percentiles. *N* > 12 cells were tested per condition. *p* values were calculated using Welch’s t-test on log_10_-transformed diffusion coefficients. ***: *p* < 0.001; **: *p* < 0.01; n.s.: *p* > 0.05. b) Chelation of divalent cations with EDTA produces a smaller increase in OM fluidity than truncation of LPS core oligosaccharides or reduction of OM proteins. Cells were grown in LB for 3 h to mid-log phase prior to EDTA treatment. Center lines indicate medians, and error bars indicate the 25^th^ and 75^th^ percentiles. *N* > 17 cells were tested per condition. *p* values were calculated using Welch’s t-test on log_10_-transformed diffusion coefficients. ***: *p* < 0.001; n.s.: *p* > 0.05.

In a previous study, we showed that divalent-cation crosslinking of LPS is critical for OM rigidity^6^. To test whether this crosslinking also restricts OM fluidity, we measured LPS diffusion in wild-type cells following treatment with high concentrations of EDTA (**Methods**). Chelation of divalent cations with EDTA concentrations up to 50 mM resulted in only minor increases in OM fluidity in either LB or MHB (**Fig. 4b and S5**), even though such concentrations reduce OM stiffness by more than 60%^6^. Supplementation with 50 mM MgCl_2_ in LB failed to restore low fluidity in Δ*bamD* cells (**Fig. S6**), further indicating that the mechanism underlying increased LPS diffusion in Δ*bamD* is largely independent of LPS crosslinking. The differential effects of EDTA treatment (**Fig. S5**) and inner core oligosaccharide truncation (**Fig. 2a**) on LPS diffusion suggest that distinct LPS intermolecular interactions control OM fluidity and OM mechanical stiffness.

### OM fluidity and stiffness are independently tunable

Our results indicate that although OM fluidity and stiffness both arise from interactions among porins, LPS, and PLs in the outer leaflet, these physical properties can be modulated independently. For instance, growing Δ*bamB* cells exhibited a 10-fold increase in OM fluidity (**Fig. S2**) while maintaining wild-type stiffness^16^. Conversely, the *imp4213* mutation and EDTA treatment each reduce OM stiffness^6^ but did not increase fluidity (**Fig. 4**).

To systematically compare these properties across perturbations, we organized fluidity and stiffness measurements from this and our previous studies^6,16^ into a phenomenological map (**Fig. 5a**). This analysis revealed distinct trends associated with perturbations affecting OM proteins and PL levels versus those altering LPS abundance or divalent-cation interactions. Notably, *mlaA\** Δ*pldA* cells retained substantial stiffness (**Fig. 5a**) despite markedly increased OM fluidity (**Fig. 1d**). Similarly, CRISPRi knockdown of *bamD* (**Table S1**) progressively increased LPS diffusion as PLs accumulated in the outer leaflet due to defective OM protein insertion, without reducing stiffness (**Fig. 5a**), consistent with the behavior of Δ*bamB* (**Fig. S2**)^16^. In contrast, perturbations that reduce LPS levels or chelate divalent cations decreased stiffness without substantially altering fluidity (**Fig. 5a**).

**Figure 5:**
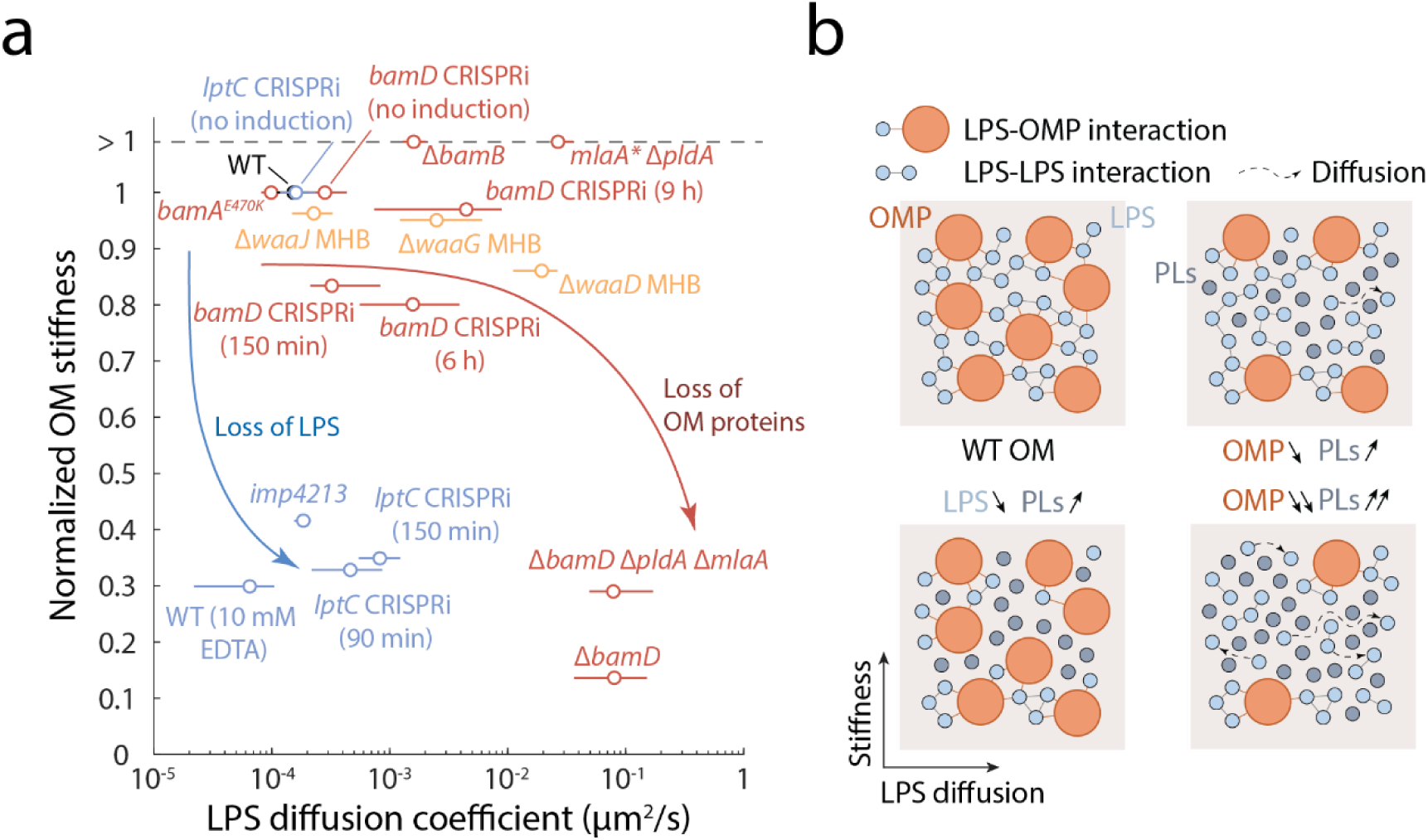
OM fluidity and stiffness can be modulated independently by perturbations to OM proteins and LPS. a) Phenomenological map comparing OM fluidity and stiffness across perturbations affecting OM composition. The *imp4213* allele, chelation of divalent cations by EDTA (which reduces LPS crosslinking and extracts LPS from the OM^77^), or knockdown of either *lpxC* or *lptC* reduced OM stiffness without substantially increasing fluidity. In contrast, Δ*bamB* cells exhibit increased fluidity while OM stiffness remains similar to wild type^16^. CRISPRi knockdown of *bamD* increased fluidity in an induction time-dependent manner under steady-state exponential growth conditions without substantially reducing stiffness. Similarly, *mlaA\** Δ*pldA* cells exhibited high fluidity while retaining near-wild-type-level stiffness. LPS core truncation mutants (*waa* mutants) grown in MHB showed increased fluidity accompanied by a modest reduction in stiffness. All fluidity measurements were obtained in this study. OM stiffness values for *imp4213*^6^, EDTA-treated wild-type^6^, *bamA^E470K^*^16^, and Δ*bamB*^16^ cells were reported previously, whereas all other stiffness measurements were performed in this study. Mutant stiffness values were normalized to their corresponding wild-type strain, parent strain, or no-induction control. Circles indicate medians, and error bars indicate the 25^th^ and 75^th^ percentiles of LPS diffusion coefficients. Error bars are not shown for stiffness, as values were calculated from averaged cell length contractions (**Methods**). b) Schematic illustrating the changes in OM composition, organization, and molecular interactions that underlie the stiffness and fluidity regimes in the corresponding quadrants of the fluidity-stiffness map in a). Reducing OM protein levels promotes PL intercalation between LPS molecules, weakening LPS packing and increasing LPS fluidity. However, PL accumulation resulting from reduced OM protein levels is insufficient to disrupt the continuity of the LPS-LPS interaction network that supports load bearing across the cell length. In contrast, replacing LPS with PLs likely disrupts this LPS interaction network and yields a more continuous OMP-PL matrix with limited load-bearing capacity, while the increased OMP to LPS ratio may still restrict LPS diffusion through steric confinement and/or LPS-OMP interactions.

Together, these results indicate that membrane composition influences OM properties through separable mechanisms, revealing two largely independent control knobs (**Fig. 5a**). Abundant LPS, stabilized by divalent cations, contributes strongly to the load-bearing capacity of the OM. When OM proteins are gradually replaced by PLs, the OM can retain near-wild-type stiffness even as PL accumulation fluidizes LPS, implying that the LPS network continues to bear most of the mechanical load as PLs incorporate into the outer leaflet. In contrast, conditions that reduce LPS abundance or disrupt cation-mediated stabilization decrease stiffness without a commensurate increase in LPS mobility. Under these conditions, porins may still restrain LPS motion within a PL-enriched outer leaflet, potentially through LPS-porin interactions.

The fluidity-stiffness map (**Fig. 5a**) also reveals distinct contributions from different structural features of LPS. Chelation of divalent cations with EDTA reduced stiffness without affecting fluidity, whereas truncation of the core oligosaccharides (*waa* mutants) progressively increased LPS mobility with only modest decreases in stiffness (**Fig. 5a**). These trends indicate that the core oligosaccharides contribute primarily to restricting lateral mobility, whereas divalent-cation-mediated salt bridges contribute more strongly to OM mechanical rigidity.

## Discussion

In this study, we used FRAP assays of fluorescently labeled LPS to quantify effective OM fluidity and identify molecular interactions that control LPS mobility. Across a wide range of genetic and chemical perturbations, PL insertion into the outer leaflet and truncation of LPS oligosaccharides markedly increased OM fluidity, whereas perturbations that reduce LPS levels or chelate divalent cations primarily weakened OM stiffness. These findings indicate that OM fluidity and stiffness can be modulated independently through distinct molecular interactions, challenging the intuition that low fluidity and high stiffness are tightly coupled and highlighting multiple molecular targets for tuning OM physical properties. Further elucidating these mechanisms will deepen our understanding of OM organization and maintenance and may ultimately reveal novel strategies for destabilizing the OM. Throughout this study, we use lateral diffusion of LPS as a readout of effective OM fluidity. Although this measurement reports specifically on LPS mobility, the central role of LPS in mediating intermolecular interactions within the membrane means that changes in LPS mobility reflect changes in the physical state of the OM surface.

Using fluidity as a readout of intermolecular interactions, our study reveals architectural features that shape OM organization. Our data suggest that when PLs replace proteins in the outer leaflet, they intercalate among LPS molecules and weaken interactions that normally constrain LPS mobility. We did not observe heterogeneous fluorescence patterns in Δ*bamD* cells (**Fig. 1c**), at least at the diffraction-limited scale, suggesting that PL accumulation does not produce obvious micron-scale segregation of LPS-rich and PL-rich regions, although we cannot exclude nanoscale domains below the diffraction limit. LPS core truncation reveals a distinct mechanism for disrupting LPS packing. In atomistic simulations, truncation reduced direct hydrogen bonding and divalent cation-mediated interactions between neighboring LPS molecules, increased local fluctuations, disrupted both local and long-range packing order, and accelerated nearest-neighbor exchange (**Fig. 3, S4**). These results suggest that intact core oligosaccharides stabilize transient local coordination shells that confine neighboring LPS molecules and limit lateral rearrangement. Core truncation weakens these coordination shells, providing a molecular explanation for the increased LPS mobility observed experimentally.

The increase in fluidity caused by reductions in LPS levels and corresponding PL accumulation (**Fig. 4a**) did not phenocopy the fluidization caused by replacement of proteins with PLs (**Fig. 1**), potentially because abundant OM proteins interact with and restrict LPS mobility or act as large obstacles that divide the membrane surface into inter-porin channels that limit diffusion (**Fig. 5b**). Consistent with this idea, OMP-LPS arrangements observed in wild-type cells may create locally confined populations of LPS alongside more mobile regions, as suggested by previous descriptions of porin-LPS organization in the OM^2,24^. Thus, disruption of LPS packing emerges as a major driver of OM fluidization, while OMPs may modulate LPS mobility through steric confinement or specific LPS-protein interactions, without being the primary determinant of LPS diffusion under the conditions tested. Consistent with this interpretation, across perturbations where OM protein levels differ substantially (*bamA^E470K^* versus Δ*bamD*), the largest shifts in LPS diffusion coincide with conditions known to alter outer-leaflet PL content. Moreover, LPS mobility is high in *mlaA*\* Δ*pldA* cells, in which the primary perturbation drives PL accumulation rather than directly disrupting OM protein assembly. Together, these observations demonstrate that increased outer-leaflet PL abundance can drive large increases in LPS mobility without directly disrupting OM protein assembly, supporting PL-mediated disruption of LPS packing as a major determinant of OM fluidization.

In contrast to fluidity, OM stiffness reflects the ability of the membrane to resist in-plane stress and, as our data suggest, is more sensitive to reductions in LPS than to reductions in OM proteins (**Fig. 5a**). We propose that maintaining stiffness requires a laterally continuous network of LPS-LPS interactions spanning the cell surface, reinforced primarily by divalent-cation bridges between neighboring LPS molecules (**Fig. 5b**). When PLs replace OM proteins, they partially disrupt local LPS packing but apparently do not fragment the LPS interaction network into mechanically disconnected regions, allowing the membrane to retain near-wild-type stiffness even as lateral mobility increases. In contrast, when PLs replace LPS, the combined decrease in LPS abundance and emergence of a more continuous OMP-PL matrix may more readily break apart the LPS interaction network. Because such an OMP-PL mixture is unlikely to sustain in-plane compression as effectively as an LPS-rich network, stiffness decreases even when lateral mobility remains constrained.

The decoupling of OM fluidity and stiffness has important implications for the structural and functional versatility of Gram-negative bacteria. Our findings suggest that while a rigid, crosslinked LPS network primarily provides mechanical strength, shifts in the balance between OM proteins and PLs modulate lateral mobility. This separation creates opportunities for bacteria to tune OM physical properties to meet diverse functional demands and environmental conditions. It is therefore likely that regulatory pathways adjust OM composition to maintain appropriate mechanical and diffusive properties. Measuring these properties across varying growth conditions, including growth phases, growth rate, and temperature, may reveal mechanisms of envelope homeostasis. In addition, structural variation in the OM across species may produce distinct mechanical and diffusive behaviors. For example, *Vibrio cholerae* has a much weaker OM than the load-bearing OM of *E. coli* or *Pseudomonas aeruginosa*^6^, raising the possibility that OM fluidity also varies across species according to ecological niche.

The OM contains abundant structural porins but relatively few enzymes, possibly reflecting constraints imposed by low lateral fluidity on surface-embedded catalysis. Nevertheless, several OM-associated enzymes could respond to and potentially regulate OM fluidity. PldA hydrolyzes PLs mislocalized to the outer leaflet^57^, while PagP transfers palmitate from PLs to lipid A in response to elevated outer-leaflet PL levels^58^. Because the activities of both enzymes are linked to disrupted lipid asymmetry, the increased fluidity accompanying PL accumulation could influence their access to membrane substrates. Such coupling between OM physical state and enzymatic activity could provide a feedback mechanism that helps restore lipid asymmetry and maintain envelope homeostasis.

## Methods

### Culture conditions for LPS staining

*E. coli* cells were grown overnight at 37 °C in M9 minimal medium supplemented with 0.2% glucose (Sigma-Aldrich, Cat. #G8270) and 0.5% casamino acids (RPI, Cat. #C45000). Overnight cultures were diluted 1:1000 into fresh M9 medium containing 5 mM 8-azido-3,8-dideoxy-D-manno-octulosonic acid (Kdo-N_3_, WuXi AppTec) and incubated overnight under the same conditions. The resulting saturated cultures were diluted 1:200 into lysogeny broth (LB, RPI, Cat. #L24060) or Mueller Hinton Broth (MHB, BD Difco, Cat. #212322) supplemented with 5 mM Kdo-N_3_ and grown for the indicated durations before LPS staining and FRAP measurements. For Δ*bamD* strains, cultures were diluted 1:50 due to the lower optical density (OD) reached in saturated cultures.

### LPS labeling via azide–alkyne click chemistry

LPS molecules were labeled using a copper-catalyzed azide–alkyne cycloaddition

reaction adapted from a published protocol^37^. A catalyst buffer was prepared by mixing 2 mM CuSO_4_ (Sigma-Aldrich, Cat. #31293) and 4 mM tris(6-galactosyltriazomethyl)amine (TGTA, WuXi AppTec)^61^ in 50 mM sodium phosphate (Sigma-Aldrich, Cat. #8210-OP) buffer (pH 7.5) and incubating overnight at 37 °C with shaking. To prepare the labeling mixture, the catalyst buffer was supplemented with 4 mM aminoguanidine HCl (Sigma-Aldrich, Cat. #396494), 5 mM sodium ascorbate, and 130 µM rhodamine 110-PEG4-alkyne (Sigma-Aldrich, Cat. #761621). Cells cultured in the presence of Kdo-N_3_ were washed three times with 50 mM sodium phosphate buffer, incubated with the labeling mixture for 10 min at room temperature, and then washed twice with sodium phosphate buffer before resuspension in the same buffer for FRAP measurements.

### Fluorescence recovery after photobleaching (FRAP) assays

One microliter of cells with fluorescently labeled LPS was placed onto a 1% agarose pad prepared with 0.85X PBS (Gibco, Cat. #70011044), which has approximately the same osmolarity as LB medium. Samples were imaged using a Zeiss LSM880 inverted laser-scanning confocal microscope equipped with a 63X oil-immersion objective. A manually defined region of the cell, typically near midcell, was photobleached by repeated high-intensity laser scanning. Fluorescence recovery within the bleached region was then monitored for 30 s to 3 min, depending on the recovery kinetics. Images were acquired using Zeiss ZEN software. All imaging was performed at 37 °C using an environmental chamber with active temperature control.

### FRAP analysis

Time-lapse fluorescence images were segmented using the software *Morphometrics*^62^ to extract cell contours at sub-pixel resolution. These contours were processed using the meshing algorithm from *MicrobeTracker*^63^ and MATLAB (R2025b) to determine the cell centerline along the major axis and generate a one-dimensional fluorescence intensity profile along the cell length. To quantify membrane fluidity, these fluorescence recovery profiles were compared to simulated one-dimensional diffusion curves with zero-flux boundary conditions, and the diffusion coefficient was extracted via linear fitting (**Fig. 1a**).

For each cell, the dynamics of the one-dimensional fluorescence profile *F*(*x*, *t*) for 0 ≤ *x* ≤ *L*, where *L* denotes cell length, were simulated using the diffusion equation with zero-flux boundary conditions:

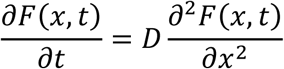

with initial condition

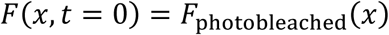

and boundary conditions

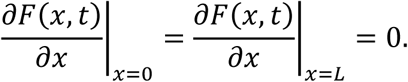

The diffusion equation was solved using the implicit backward Euler method.

The progress of fluorescence recovery was quantified using a normalized recovery metric *R*(*t*), where 0 < *R*(*t*) < 1. This metric was defined as the normalized L_1_-norm difference between the fluorescence profile at time *t* and the profile corresponding to complete equilibration of labeled molecules after long diffusion times (**Fig. S1a and S3a**), i.e.,

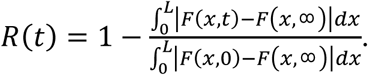

For linear diffusion, *R*(*t*) is predicted to converge approximately exponentially from 0 to 1. For FRAP experiments, the long-time profile *F*_exp_(*x*, ∞) was approximated by the pre-photobleaching fluorescence profile corrected for total fluorescence loss due to photobleaching. For simulations, the long-time profile *F*_sim_(*x*, ∞) was defined as uniform with the same total fluorescence as the post-photobleaching profile.

For each cell, experimental FRAP recovery profiles were first normalized to the total fluorescence immediately after photobleaching and then compared to a series of simulated profiles with sufficient temporal resolution to densely cover recovery values *R*_sim_(*t*) from 0 to 0.99. Simulations were performed using a standard diffusion coefficient of *D*_sim_ = 1 µm^2^/s and a normalized length of 1 µm (the cell length was later scaled to the experimentally measured value).

The difference between each experimental and simulated profile was calculated as

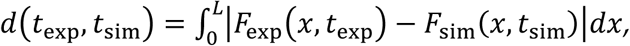

which was then normalized to obtain a similarity matrix *E*(*t*_exp_, *t*_sim_):

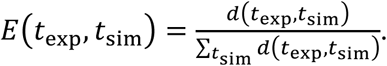

To determine the experimental diffusion coefficient, a linear regression was performed using all (*t*_exp_, *t*_sim_) data pairs weighted by *E*(*t*_exp_, *t*_sim_). The resulting slope *p* was then used to scale the standard diffusion coefficient *D*_sim_ to the experimental coefficient as

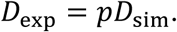

Because this method relies on comparing fluorescence recovery between experiments and simulations, it effectively covers diffusion rates above 10^-4^ µm^2^/s. For slower diffusion, corresponding to characteristic timescales greater than 10^4^ s, fluorescence recovery is negligible within the observation window (∼2-3 min), resulting in low signal-to-noise ratios and inaccurate estimates of diffusion coefficients. In such cases, LPS fluidity is effectively immobile relative to physiologically relevant timescales such as the 20-60 min doubling time in standard culture media.

To assess deviations from ideal linear diffusion, we compared experimental recovery *R*_exp_(*t*_exp_) with simulated recovery *R*_sim_(*t*_exp_; *D*_exp_) for each cell (**Fig. S1b and S3b**). Cells with data points falling along the diagonal of the *R*_exp_-*R*_sim_ plot exhibited kinetics consistent with linear diffusion, whereas cells with convex deviations toward the *R*_exp_ axis showed progressively slowing recovery, potentially reflecting subpopulations of less mobile LPS molecules.

To estimate the fraction of immobile LPS in each cell, the measured recovery curve *R*_exp_(*t*_exp_) was fitted to

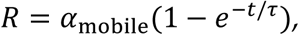

where *α*_mobile_ and *τ* represent the fraction of mobile LPS and the characteristic recovery timescale, respectively (**Fig. S1c and S3c**). This analysis assumes that the mobile LPS population follows approximately normal diffusion, which was largely supported under the conditions tested in this study (**Fig. S1 and S3**).

### OM stiffness measurements

Stiffness was quantified using a microfluidic plasmolysis-lysis assay as previously described^6^. Steady-state log-phase cells grown in LB or MHB with 300 µM 3-[[(7-hydroxy-2-oxo-2H-1-benzopyran-3-yl)carbonyl]amino]-D-alanine hydrochloride (HADA, MedChemExpress, Cat. #HY-131045/CS-0124027) were loaded into a CellASIC ONIX microfluidic flow cell (Sigma-Aldrich, Cat. #B04A-03-5PK) and incubated for 30 min at 37 °C in the corresponding medium containing HADA. Cells were then perfused with 0.85X PBS, subjected to a hyperosmotic shock with 1 M sorbitol (Sigma-Aldrich, Cat. #S6021) in 0.85X PBS, and subsequently treated with 5% (w/v) N-lauroylsarcosine sodium salt (MP biomedicals, Cat. #190289) in 1 M sorbitol and 0.85X PBS as a detergent to remove the membranes. Media exchanges were performed using the CellASIC ONIX2 microfluidic platform (Sigma-Aldrich, Cat. #CAX2-S0000). Cells were imaged using a Nikon Eclipse Ti-E inverted fluorescence microscope equipped with a 100X oil-immersion objective (NA 1.40; Nikon Instruments). Phase-contrast and fluorescence images were acquired with a Prime BSI Express sCMOS camera (Teledyne Photometrics) to monitor cell morphology. All imaging was conducted at 37 °C using an environmental chamber with active temperature control (HaisonTech).

Images of HADA-labeled cells were segmented using the machine-learning-based software *DeepCell*^64^ and analyzed with *Morphometrics* to extract single-cell contours. Because HADA is sensitive to photobleaching, images were acquired at a relatively low frame rate (one frame every 1–5 min). Cell lineages were therefore reconstructed by manual curation for a representative subset of cells. Cell length and width were calculated from the contours using the *MicrobeTracker* meshing algorithm.

Cell wall lengths were measured in the turgid state (*l*_1_), under plasmolysis (*l*_2_), and after OM removal (*l*_3_). Length contractions between these states were calculated as ε_12_ = (*l*_1_−*l*_2_)/*l*_2_, ε_23_ = (*l*_2_−*l*_3_)/*l*_3_, and ε_13_ = (*l*_1_−*l*_3_)/*l*_3_. The OM stiffness *k*_OM_ relative to the cell wall stiffness *k*_CW_ was estimated using the average length contractions according to *k*_OM_ = *k*_CW_ <ε_23_>/[<ε_12_> (1 + <ε_23_>)], where <·> denotes the mean of the indicated quantity, as previously described^6^.

### CRISPR knockdown

EZ rich defined medium (EZ-RDM) was prepared by combining 100 mL of 10X MOPS buffer (Teknova, Cat. #M2101), 100 mL of 10X ACGU solution (Teknova, Cat. #M2103), 10 mL of 0.132 M K_2_HPO_4_ (Sigma-Aldrich, Cat. #PX1570), and 200 mL of 5X supplemented EZ (Teknova, Cat. #M2104) with 580 mL of water^65^. The medium was supplemented with 0.2% glycerol (Sigma-Aldrich, Cat. #GX0190).

*lpxC* and *lptC* CRISPRi strains were cultured and induced following a previously established protocol^66^ with minor modifications for LPS labeling. For FRAP assays, cells were grown overnight at 37 °C in M9 minimal medium, diluted 1:1000 into fresh M9 medium containing 5 mM Kdo-N_3_, and grown overnight under the same conditions. The resulting culture was diluted 1:200 into EZ-RDM supplemented with 0.2% glycerol and 5 mM Kdo-N_3_ and grown for 3 h to mid-log phase. Knockdown was induced by adding 0.2% L-arabinose (Sigma-Aldrich, Cat. #A91906) and cells were harvested after 90 or 150 min for FRAP measurements.

For OM stiffness measurements, cells were grown overnight at 37 °C in EZ-RDM supplemented with 0.2% glycerol, diluted 1:200 into fresh EZ-RDM, and grown for 3 h before being loaded into a CellASIC flow cell. In the flow cell, knockdown was induced with 0.2% L-arabinose in EZ-RDM for 90 or 150 min before plasmolysis with 1 M sorbitol in 0.85X PBS and lysis with 5% N-lauroylsarcosine sodium salt. For these experiments, 25 µg/mL wheat germ agglutinin (WGA) Alexa Fluor 488 (Invitrogen, Cat. #W11261), rather than HADA, was used throughout the experiment to achieve improved cell envelope staining.

The *bamD* CRISPRi strain was obtained from a previously generated CRISPR interference library^67^ and cultured and induced according to the established protocol^67^ with minor modifications for LPS labeling. For FRAP assays, cells were grown overnight at 37 °C in M9 minimal medium, diluted 1:1000 into fresh LB containing 5 mM Kdo-N_3_, and grown overnight under the same conditions. The resulting culture was diluted 1:200 into LB containing 5 mM Kdo-N_3_ and grown for 3 h to mid-log phase. Knockdown was induced by adding IPTG to a final concentration of 1 mM, and cultures were maintained in exponential growth by serial dilution for 9 h. Cells were harvested for FRAP measurements at 0 min, 150 min, 6 h, and 9 h after induction.

### Molecular dynamics simulations

All-atom models of the *E. coli* K-12 OM were constructed using CHARMM-GUI^68,69^. The native membrane composition consisted of an outer leaflet containing 100% homogeneous LPS molecules and an inner leaflet composed of 90% PVPE, 5% PVPG, and 5% PVCL2 lipids^70,71^. Each membrane patch contained 34 LPS molecules in the outer leaflet. Each system was solvated in 0.15 M KCl, and LPS molecules were neutralized with Mg^²2^ and Ca^²2^ ions. Simulations were performed using the CHARMM36 all-atom force field^72^ and NAMD3 molecular dynamics software^73^. Production simulations employed hydrogen mass repartitioning to enable a 4 fs integration timestep^74^. Bonds involving hydrogen were constrained. Van der Waals interactions were smoothly switched to zero between 10 and 12 Å. Long-range electrostatics were treated using the particle mesh Ewald method with a grid spacing of 1.0 Å. Simulations were carried out under periodic boundary conditions at 310 K and 1 atm using Langevin dynamics with a damping coefficient of 1 ps⁻¹ and the Langevin piston method. Semi-isotropic pressure coupling was used to allow independent fluctuations of the membrane plane and bilayer normal. Two independent replicas were simulated for each system, yielding two wild-type trajectories and two Δ*waaF* trajectories. Trajectory coordinates were saved every 0.2 ns for analysis. Each system was simulated for 5 µs. Before analysis, global translation in the membrane plane was removed by aligning each trajectory to the combined center of mass of all LPS molecules.

Trajectory processing and analysis were performed using custom Tcl and Python scripts together with VMD^75^. Nearest-neighbor calculations were performed in Python using SciPy^76^, and figures were generated using Python.

Hydrogen bonds between LPS molecules were analyzed using VMD. For each LPS molecule, neighboring LPS molecules were identified by selecting molecules containing at least one atom within 4 Å of the reference LPS. Hydrogen bonds were then detected using a donor–acceptor distance cutoff of 3.5 Å and a donor–hydrogen–acceptor angle cutoff of 30°. Both donor–acceptor directions were evaluated to capture hydrogen bonds in which the reference LPS acted as either donor or acceptor. Hydrogen-bond counts were averaged over the final 1 µs of each trajectory (**Fig. 3a,b**).

To estimate lateral diffusion coefficients, we calculated the two-dimensional mean-squared displacement (MSD) of the LPS center of mass in the membrane plane. MSD was calculated separately for five consecutive 1-µs intervals within each trajectory, with the reference coordinates reset at the beginning of each interval at 0, 1, 2, 3, and 4 µs. For each interval, the MSD was averaged over all 34 LPS molecules and fitted using a linear function. The diffusion coefficient was calculated as one-quarter of the fitted MSD slope, assuming two-dimensional diffusion. Because the trajectories primarily sampled local motion within a confined regime rather than long-range diffusion, these values should be interpreted as effective diffusion estimates over the simulated timescale (**Fig. S4b,c**).

To quantify local LPS mobility, we calculated the root-mean-square fluctuation (RMSF) of the lipid A center of mass in the membrane plane during the final 1 µs of each trajectory. Global translation was removed using the combined center of mass of all LPS molecules. RMSF was calculated separately for each LPS molecule and compared between wild-type and Δ*waaF* systems (**Fig. 3e,f**).

To quantify LPS packing, we calculated local and global sixfold bond-orientational order parameters from the two-dimensional lipid A center-of-mass positions. Periodic boundaries in the membrane plane were accounted for when calculating intermolecular distances. For each LPS molecule i, the six nearest neighboring LPS molecules were identified, and the hexatic order parameter was calculated as

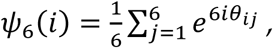

where *θᵢⱼ* is the angle between the vector connecting LPS *i* to neighbor *j* and a fixed axis in the membrane plane. The magnitude |*ψ*_6_(*i*)| ranges from 0 to 1, with values approaching 1 indicating that the six neighbors are arranged close to ideal hexagonal symmetry. Mean local order was calculated by averaging |*ψ*_6_(*i*)| over all LPS molecules in each frame:

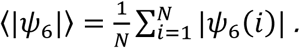

For per-LPS comparisons, local order was averaged over frames during the final 1 µs of each trajectory for each LPS molecule, yielding the time-averaged local order 〈|*ψ*_6_(*i*)|〉*_t_* (**Fig. 3h**). Global hexatic order was calculated as

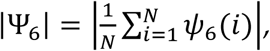

where *N* is the number of LPS molecules. In contrast to mean local order, which measures the degree of hexagonal organization within individual neighborhoods, global order is sensitive to common orientational alignment across the entire LPS layer. Time courses were calculated over the full trajectories, and distributions used for comparison were obtained from the final 1 µs of each simulation (**Fig. 3g-i, S4d**).

To quantify the persistence of local LPS coordination, we calculated a nearest-neighbor retention function from the two-dimensional lipid A center-of-mass positions. Periodic boundaries in the membrane plane were included in all nearest-neighbor searches. For each LPS molecule *i*, the six nearest neighbors at the first analyzed frame were defined as the initial neighbor set *N*ᵢ(0). At each subsequent time point, the six nearest neighbors were recalculated to obtain *N*ᵢ(t). The fraction of initial neighbors retained at time *t* was calculated as

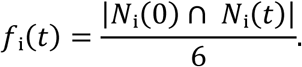

The neighbor-retention function was obtained by averaging fᵢ(t) over all LPS molecules. A value of 1 indicates complete preservation of the initial coordination shell, whereas decreasing values indicate progressive neighbor exchange and lateral rearrangement (**Fig. 3j**).

Mean interaction energies were calculated using the NAMD Energy plugin, considering the electrostatic and van der Waals interactions between each LPS molecule and its nearest neighbors (**Supplementary Movies S1–S4**). In this analysis, each LPS was treated as a distinct molecular unit, including any divalent cations within 4 Å due to their role in stabilizing intermolecular interactions. Because hydrogen bonds are not treated as a separate term in this calculation, their contributions are implicitly included within the electrostatic component.

Similarly, pairwise interaction energies were calculated between each LPS molecule and every other LPS in the membrane, again including nearby divalent cations within 4 Å in each LPS selection. These data were used to construct time-resolved inter-LPS interaction networks (**Supplementary Movies S5–S8**).

To quantify divalent-cation association with LPS, we counted Mg^²2^ and Ca^²2^ ions located within 4 Å of each LPS molecule in every analyzed frame during the final 1 µs of each trajectory. Counts were averaged over time separately for each LPS molecule, producing one mean cation-association value per LPS molecule for comparison between systems (**Fig. 3c,d**).

## Data and code availability

All data used for generating figures in this study are available at a Stanford Digital Repository: https://purl.stanford.edu/gg071dr7283.

## Supplementary Figures

**Figure S1:**
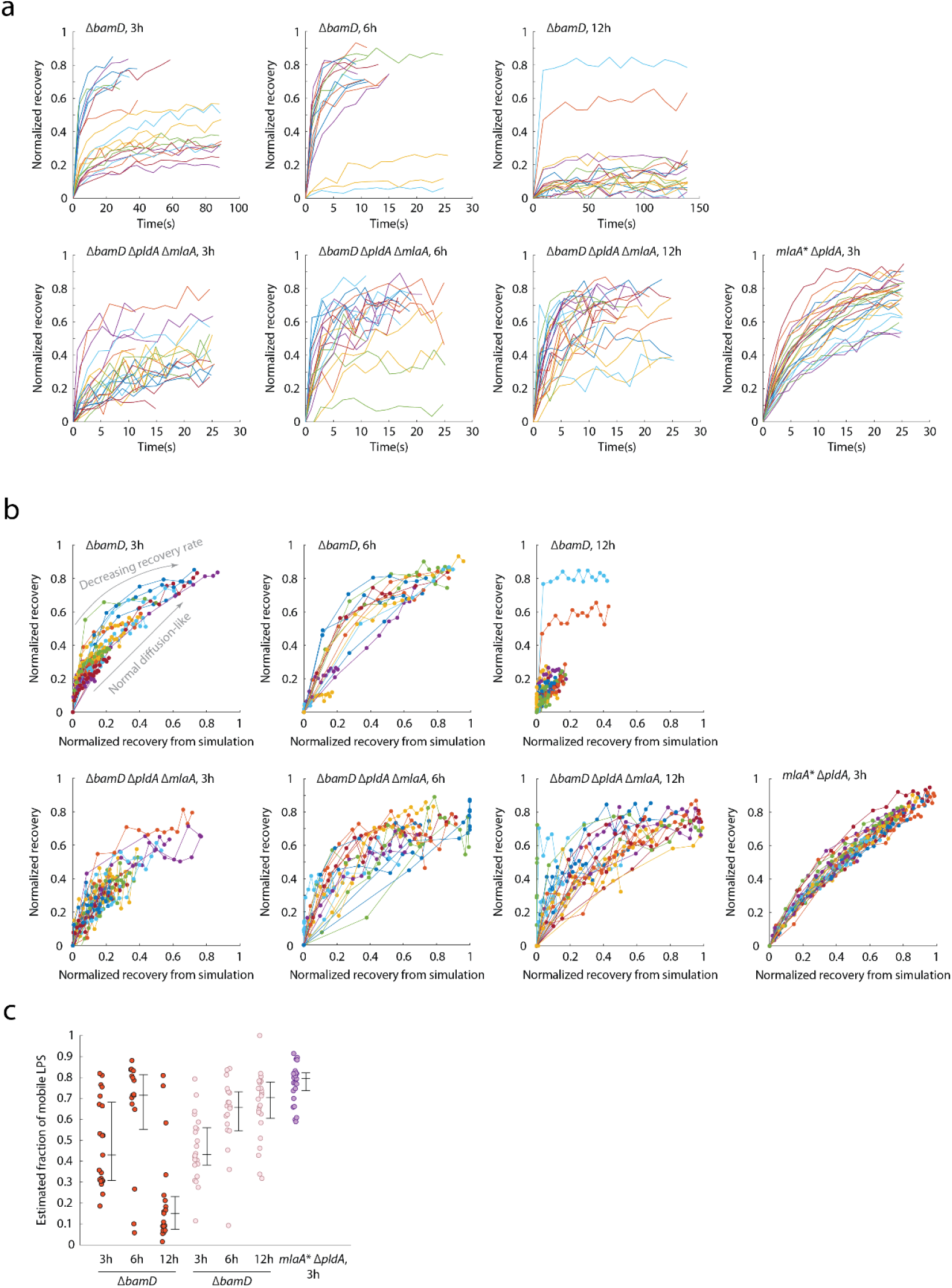
Fluorescence recovery of labeled LPS largely resembles normal diffusion in mutants with increased PL content in the OM outer leaflet. Across these conditions, recovery kinetics were broadly consistent with normal diffusion along the cell’s long axis (**Methods**), with minor deviations arising from a gradual decrease in recovery rate in some cells, suggesting subpopulations of more confined LPS molecules. Nonetheless, this confinement did not halt LPS motion, as indicated by the low estimated fraction of immobile LPS (**Methods**). Although ensemble FRAP measurements do not necessarily reflect Brownian motion at the molecular scale, the limited deviation from behavior predicted by linear diffusion suggests that PL insertion globally mobilizes LPS across the OM. a) Normalized fluorescence recovery (**Methods**) of individual cells with reduced OM protein levels and/or increased PL content, measured across genetic perturbations and growth phases. Δ*bamD* (3 and 6 h), Δ*bamD* Δ*pldA* Δ*mlaA* (3, 6, and 12 h), and *mlaA*\* Δ*pldA* (3 h) cells exhibit rapid fluorescence recovery after photobleaching. Values of 0 and 1 represent the start and long-timescale limit of recovery, respectively, with *t*=0 marking the time point immediately after photobleaching. b) Comparison between simulated and experimental normalized recovery. Δ*bamD* (3 and 6 h), Δ*bamD* Δ*pldA* Δ*mlaA* (3, 6, and 12 h), and *mlaA*\* Δ*pldA* (3 h) cells exhibit only minor deviations from the line *y*=*x*, indicating recovery kinetics resembling normal diffusion. c) Mutant cells with high OM fluidity also exhibit low fractions of immobile LPS, as estimated from the normalized recovery curves (**Methods**). Center lines indicate medians, and error bars indicate the 25^th^ and 75^th^ percentiles.

**Figure S2:**
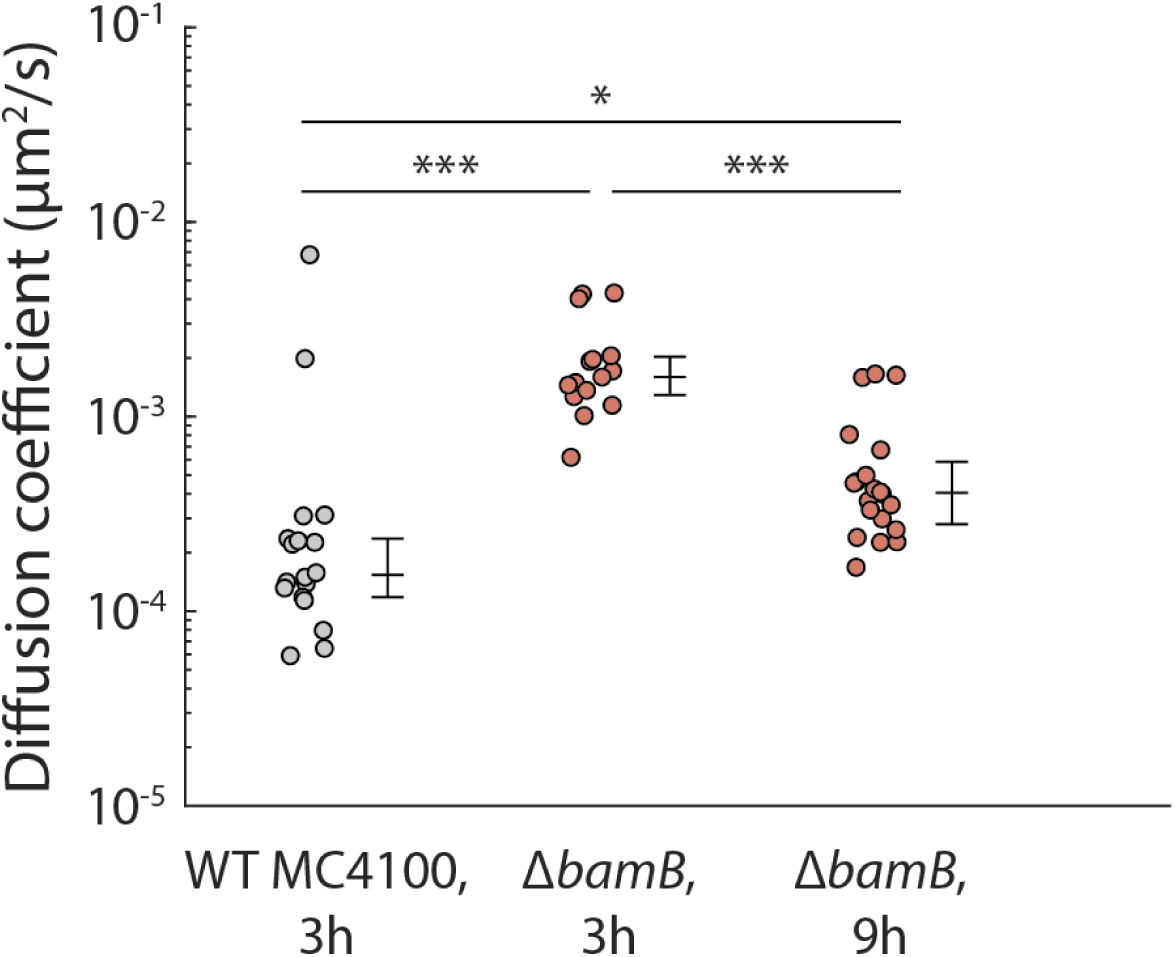
Insertion of PLs into the outer leaflet in Δ*bamB* cells increases OM fluidity. The increase in fluidity observed in Δ*bamB* cells grown in LB was growth-phase dependent and was lower than that observed in Δ*bamD* cells (Fig. 1b). Center lines indicate medians, and error bars indicate the 25^th^ and 75^th^ percentiles. *N* > 14 cells per condition. *p* values were calculated using Welch’s t-test on log_10_-transformed diffusion coefficients. ***: *p* < 0.001; *: *p* < 0.05.

**Figure S3:**
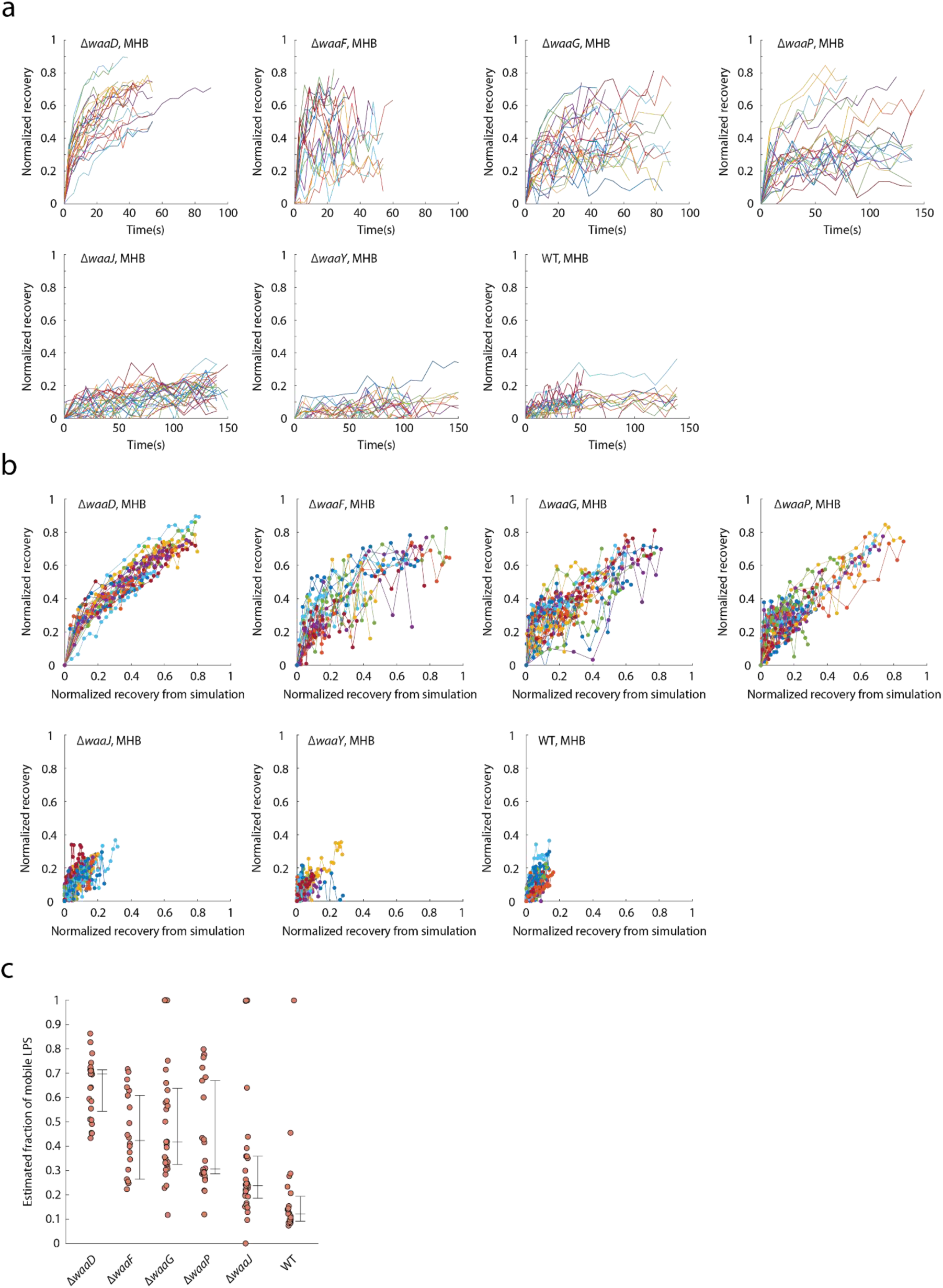
LPS core truncation increases OM fluidity and alters fluorescence recovery dynamics. a) Normalized fluorescence recovery curves (**Methods**) for individual *waa* mutant cells with truncated LPS. Values of 0 and 1 represent the start and long-timescale limit of recovery, respectively, with *t*=0 corresponding to the time point immediately after photobleaching. b) Comparison between experimentally measured fluorescence recovery and simulated recovery assuming normal diffusion. Mutants with truncated LPS show recovery dynamics that largely follow the *y*=*x* relationship across *waa* deletions. c) Fraction of mobile LPS as a function of the remaining length of the core oligosaccharide. Shorter LPS cores are associated with higher OM fluidity and an increased mobile fraction. Center lines indicate medians, and error bars indicate the 25^th^ and 75^th^ percentiles.

**Figure S4:**
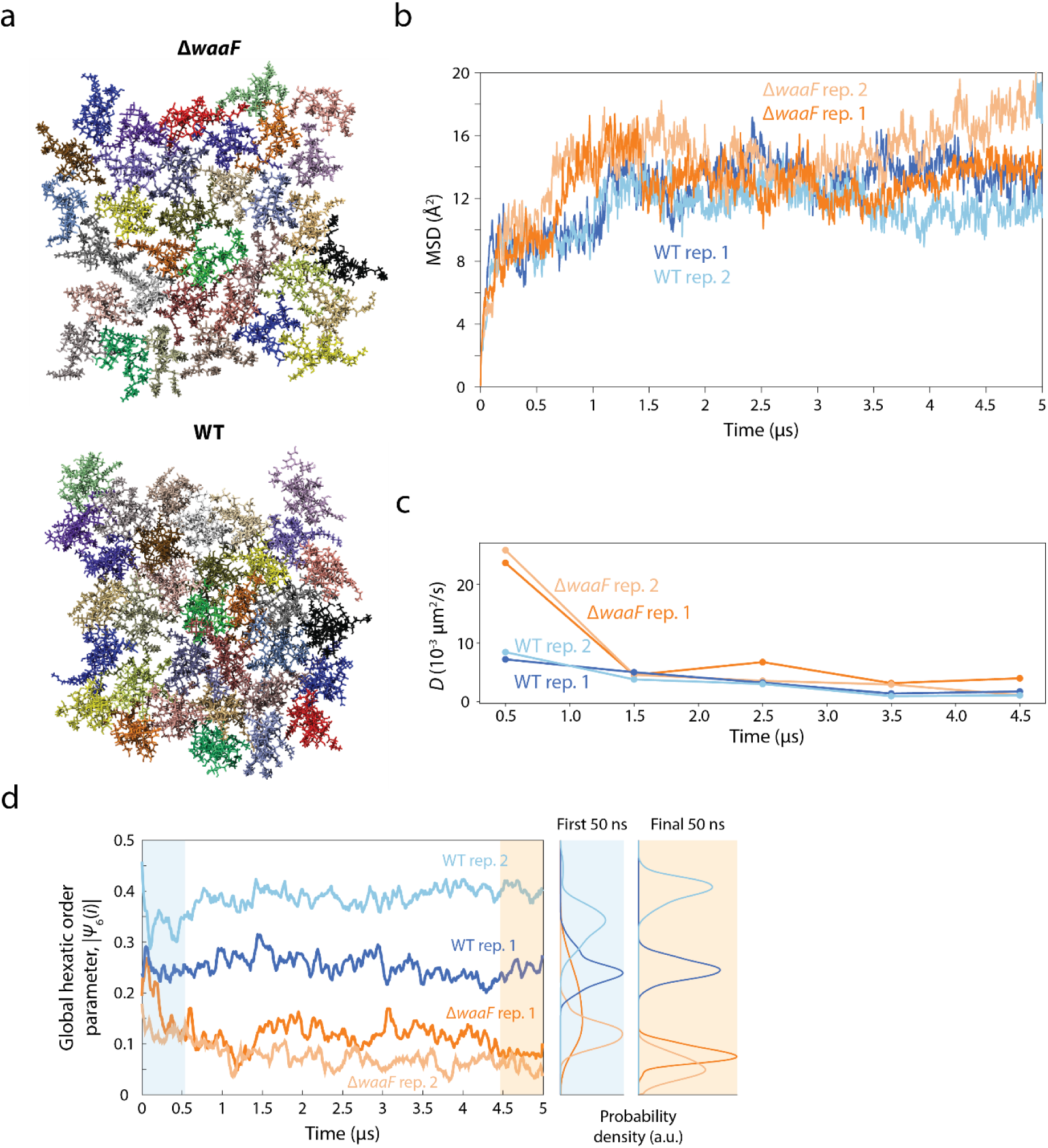
MD simulations yield estimates of LPS diffusion coefficients consistent with experimental measurements. a) Snapshots of all-atom MD simulations of OM patches containing 34 LPS molecules from wild-type or Δ*waaF* systems (**Methods**). b) Mean squared displacement (MSD) of LPS molecules during the simulations. Δ*waaF* LPS exhibits a faster initial increase in MSD during the first ∼1 µs, followed by a plateau as molecules become locally confined by neighboring LPS. Because the total simulation time (5 µs) is much shorter than the estimated time (∼1 ms) required for an LPS molecule to diffuse across its own lateral dimension, the simulations only capture local motion within a confinement regime. c) Diffusion coefficients estimated from MSD over 1 µs intervals (D∼10^-3^ µm^2^/s) are broadly consistent with values measured by FRAP (D∼10^-3^-10^-4^ µm^2^/s). d) Wild-type and Δ*waaF* LPS relax into distinct global packing states, quantified by a global hexatic order parameter, |*Ψ*_6_|, that reports the global organization of nearest neighbors (**Methods**). Time courses of |*Ψ*_6_| show that truncated LPS adopts a less ordered global packing state than wild-type LPS across independent simulations.

**Figure S5:**
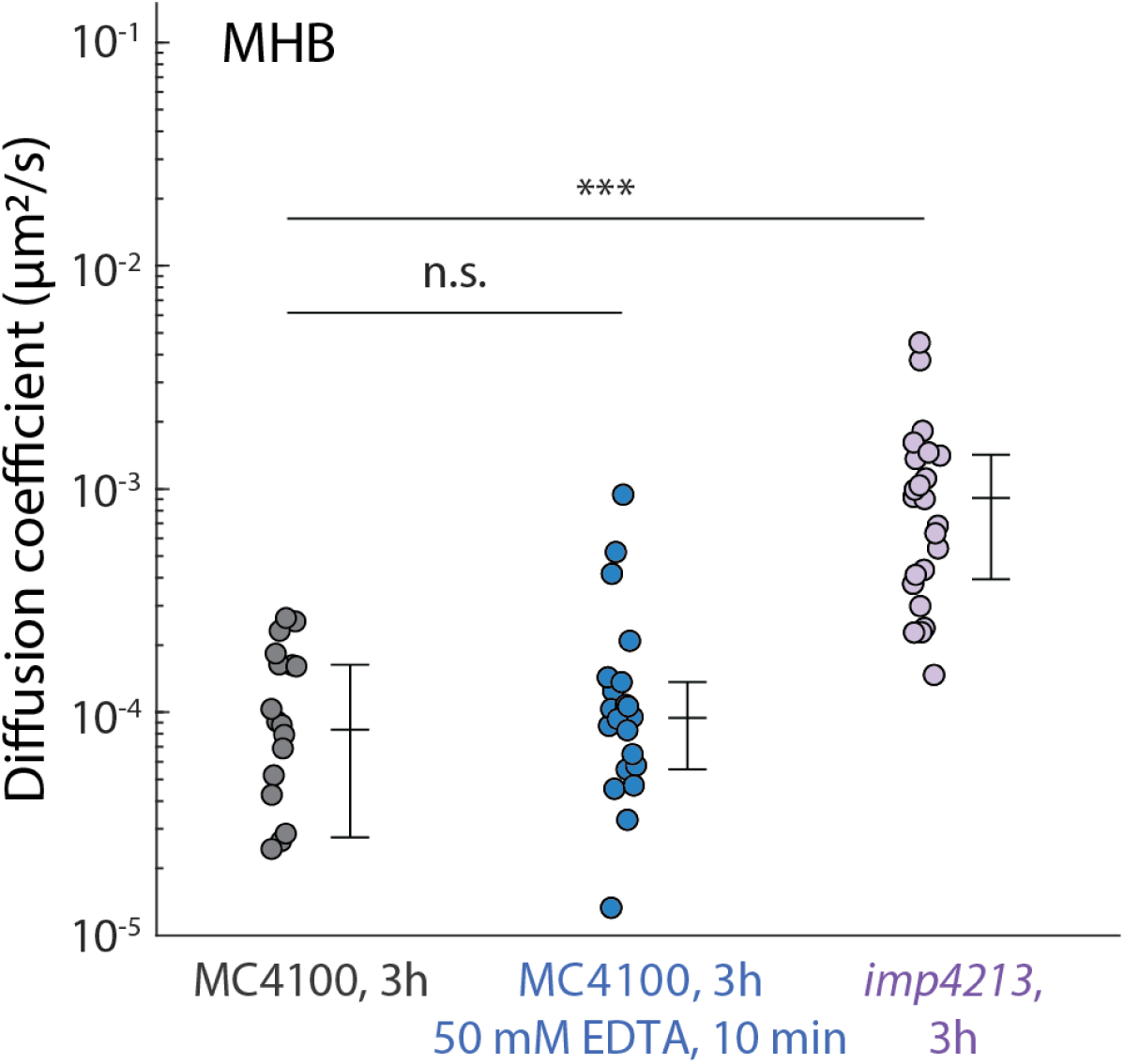
Disrupting LPS crosslinking or reducing LPS levels causes only modest increases in OM fluidity in MHB. Treating wild-type MC4100 cells grown in MHB for 3 h with 50 mM EDTA for 10 min did not significantly increase OM fluidity, consistent with results obtained in LB (Fig. 4b). The *imp4213* mutation increased fluidity by ∼10-fold in MHB, a larger effect than observed in LB (Fig. 4a), but still substantially lower than the fluidization caused by phospholipid accumulation in the outer leaflet (Fig. 1). Center lines indicate medians, and error bars indicate the 25^th^ and 75^th^ percentiles. *N* > 19 cells per condition. *p* values were calculated using Welch’s t-test on log_10_-transformed diffusion coefficients. ***: *p* < 0.001; n.s.: *p* > 0.05.

**Figure S6:**
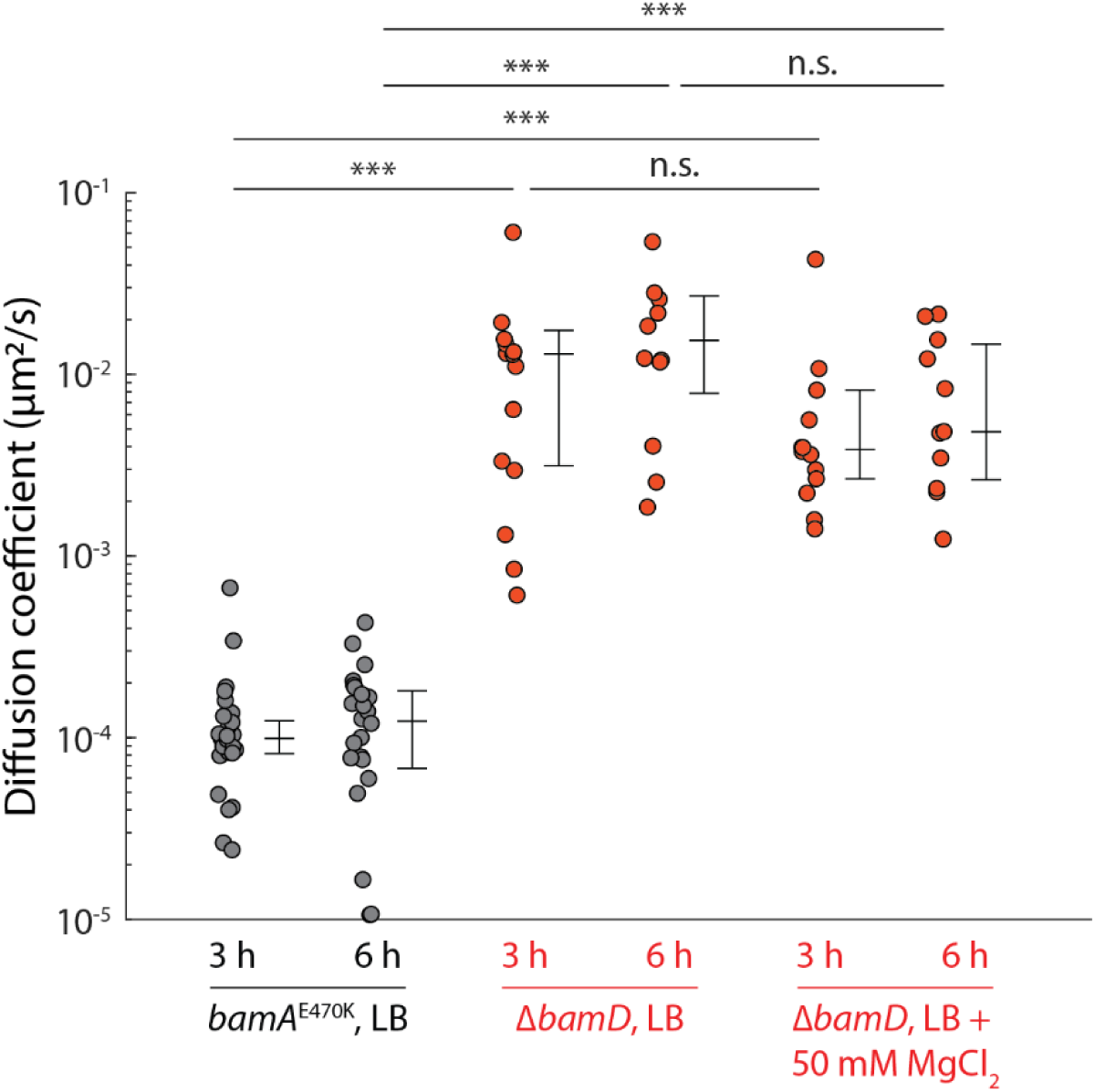
Log-phase Δ*bamD* cells maintain high OM fluidity despite Mg^2+^ supplementation. Addition of 50 mM MgCl₂ to LB did not substantially reduce OM fluidity in Δ*bamD* cells during exponential growth. Cells were grown in LB for 3 h or 6 h prior to measurement. Center lines indicate medians, and error bars indicate the 25^th^ and 75^th^ percentiles. *N* > 10 cells per condition. *p* values were calculated using Welch’s t-test on log_10_-transformed diffusion coefficients. ***: *p* < 0.001; n.s.: *p* > 0.05.

## Supplementary Movies

**Movie S1: Visualization of LPS interaction energies during an MD simulation of an *E. coli* OM patch of wild-type LPS (replicate 1).** The movie shows the nonbonded interaction energy between each of the 34 individual LPS molecules and their nearest neighbors. In the calculations, each LPS molecule was treated as a separate unit and included divalent cations (Mg^2+^, Ca^2+^) within 4 Å to account for their role in stabilizing intermolecular interactions.

**Movie S2: Visualization of LPS interaction energies during an MD simulation of an *E. coli* OM patch of wild-type LPS (replicate 2).** Visualization and interaction-energy calculations are as described for Movie S1.

**Movie S3: Visualization of LPS interaction energies during an MD simulation of an *E. coli* OM patch of Δ*waaF* LPS (replicate 1).** Visualization and interaction-energy calculations are as described for Movie S1.

**Movie S4: Visualization of LPS interaction energies during an MD simulation of an *E. coli* OM patch of Δ*waaF* LPS (replicate 2).** Visualization and interaction-energy calculations are as described for Movie S1.

**Movie S5: Time evolution of inter-LPS interaction networks during an MD simulation of an *E. coli* OM patch of wild-type LPS (replicate 1).** Each LPS molecule is represented by a sphere at its center of mass, with cylinders connecting interacting pairs. Cylinder thickness is proportional to the magnitude of the pairwise nonbonded interaction energy (scaled by 0.001), while blue and red indicate favorable (negative) and unfavorable (positive) interactions. The central simulation cell and its eight neighboring periodic images are shown to visualize interactions across periodic boundaries; non-hydrogen atoms of LPS molecules are shown in the central cell for structural context. Pairwise interaction energies were calculated using the `namdenergy` plugin with periodic electrostatics, with each LPS selection including Ca^²2^/Mg^²2^ ions within 4 Å.

**Movie S6: Time evolution of inter-LPS interaction networks during an MD simulation of an *E. coli* OM patch of wild-type LPS (replicate 2).** Visualization and interaction-energy calculations are as described for Movie S5.

**Movie S7: Time evolution of inter-LPS interaction networks during an MD simulation of an *E. coli* OM patch of Δ*waaF* LPS (replicate 1).** Visualization and interaction-energy calculations are as described for Movie S5.

**Movie S8: Time evolution of inter-LPS interaction networks during an MD simulation of an *E. coli* OM patch of Δ*waaF* LPS (replicate 2).** Visualization and interaction-energy calculations are as described for Movie S5.

## Supplementary Table

**Table S1.** *E. coli* strains used in this study.

| Strain | Genotype | Source/reference |
| --- | --- | --- |
| KC671 | MC4100 *ara*<sup>+</sup> | 78 |
| IMB1047 | MC4100 *ara*<sup>R</sup> *bamA*<sup>E470K</sup> | 16 |
| IMB1056 | MC4100 *ara*<sup>R</sup> *bamA*<sup>E470K</sup> Δ*bamD* | 16 |
| IMB1132 | MC4100 *ara*<sup>R</sup> *bamA*<sup>E470K</sup> Δ*bamD* Δ*pldA* Δ*mlaA* | 16 |
| HC726 | MC4100 *ara*<sup>+</sup> Δ*yfdI* *mlaA*<sup>*</sup> *pldA::kan* | 46 |
| BH92 | MC4100 *ara*<sup>R</sup> *bamB::kan* | 16 |
| BW25113 | $\Delta$(*araD-araB*)567 $\Delta$(*rhaD-rhaB*)568 Δ*lacZ*4787<br>(::*rrnB*-3) *hsdR*514 *rph*-1 | Gift from Jie Xiao |
| Δ*waaD* | BW25113 Δ*waaD::kan* | 51 |
| Δ*waaF* | BW25113 Δ*waaF::kan* | 51 |
| Δ*waaG* | BW25113 Δ*waaG::kan* | 51 |
| Δ*waaP* | BW25113 Δ*waaP::kan* | 51 |
| Δ*waaJ* | BW25113 Δ*waaJ::kan* | 51 |
| Δ*waaY* | BW25113 Δ*waaY::kan* | 51 |
| *lptC*<br>CRISPRi | SJ_XTL219 *lptC* sgRNA | 66 |
| *lpxC*<br>CRISPRi | SJ_XTL219 *lpxC* sgRNA | 66 |
| *bamD*<br>CRISPRi | BW25113 Tn7att::Tn7C59,<br>λatt::sgRNA(*bamD*) | 67 |
| *imp4213* | MC4100 *imp4213* *carB::Tn10* | 44 |

## Acknowledgements

The authors thank members of the Huang lab for helpful discussions. This work was supported by a Stanford Interdisciplinary Graduate Fellowship (to J.S.), NIH Awards RM1 GM135102 (to K.C.H.) and R01 GM148586 (to J.C.G.), and NSF Award EF-2125383 (to K.C.H.). K.C.H. is a Chan Zuckerberg Biohub Investigator. This work was also supported in part by the National Science Foundation under Grant PHYS-1607611 and by the hospitality of the Aspen Center for Physics.

## References

1 Silhavy, T. J., Kahne, D. & Walker, S. The bacterial cell envelope. Cold Spring Harbor perspectives in biology 2, a000414 (2010).

2. Benn, G. et al. Phase separation in the outer membrane of Escherichia coli. Proceedings of the National Academy of Sciences 118, e2112237118 (2021).

3 Sun, J., Rutherford, S. T., Silhavy, T. J. & Huang, K. C. Physical properties of the bacterial outer membrane. Nature Reviews Microbiology 20, 236–248 (2022).

4 Delcour, A. H. Outer membrane permeability and antibiotic resistance. Biochimica et Biophysica Acta (BBA)-Proteins and Proteomics 1794, 808–816 (2009).

5 Maher, C. & Hassan, K. A. The Gram-negative permeability barrier: tipping the balance of the in and the out. MBio 14, e01205–01223 (2023).

6 Rojas, E. R. et al. The outer membrane is an essential load-bearing element in Gram-negative bacteria. Nature 559, 617–621 (2018).

7 Mathelié-Guinlet, M., Asmar, A. T., Collet, J.-F. & Dufrêne, Y. F. Lipoprotein Lpp regulates the mechanical properties of the E. coli cell envelope. Nature communications 11, 1789 (2020).

8 Samsudin, F., Ortiz-Suarez, M. L., Piggot, T. J., Bond, P. J. & Khalid, S. OmpA: a flexible clamp for bacterial cell wall attachment. Structure 24, 2227–2235 (2016).

9 Szczepaniak, J. et al. The lipoprotein Pal stabilises the bacterial outer membrane during constriction by a mobilisation-and-capture mechanism. Nature communications 11, 1305 (2020).

10 Benn, G. et al. OmpA controls order in the outer membrane and shares the mechanical load. Proceedings of the National Academy of Sciences 121, e2416426121 (2024).

11 Asmar, A. T. & Collet, J.-F. Lpp, the Braun lipoprotein, turns 50—major achievements and remaining issues. FEMS microbiology letters 365, fny199 (2018).

12 Silale, A. et al. Dual function of OmpM as outer membrane tether and nutrient uptake channel in diderm Firmicutes. Nature Communications 14, 7152 (2023).

13 Deghelt, M. et al. Peptidoglycan-outer membrane attachment generates periplasmic pressure to prevent lysis in Gram-negative bacteria. Nat Microbiol 10, 1963–1974 (2025). 10.1038/s41564-025-02058-9

14 Hummels, K. R. et al. Coordination of bacterial cell wall and outer membrane biosynthesis. Nature 615, 300–304 (2023).

15 Mamou, G. et al. Peptidoglycan maturation controls outer membrane protein assembly. Nature 606, 953–959 (2022).

16 Mikheyeva, I. V., Sun, J., Huang, K. C. & Silhavy, T. J. Mechanism of outer membrane destabilization by global reduction of protein content. Nature communications 14, 5715 (2023).

17 Fivenson, E. M. et al. A role for the Gram-negative outer membrane in bacterial shape determination. Proceedings of the National Academy of Sciences 120, e2301987120 (2023).

18 Chavent, M. et al. How nanoscale protein interactions determine the mesoscale dynamic organisation of bacterial outer membrane proteins. Nature communications 9, 2846 (2018).

19 Webby, M. N. et al. Lipids mediate supramolecular outer membrane protein assembly in bacteria. Science Advances 8, eadc9566 (2022).

20 Herrmann, M., Schneck, E., Gutsmann, T., Brandenburg, K. & Tanaka, M. Bacterial lipopolysaccharides form physically cross-linked, two-dimensional gels in the presence of divalent cations. Soft matter 11, 6037–6044 (2015).

21 Ursell, T. S., Trepagnier, E. H., Huang, K. C. & Theriot, J. A. Analysis of surface protein expression reveals the growth pattern of the gram-negative outer membrane. (2012).

22 Lithgow, T., Stubenrauch, C. J. & Stumpf, M. P. Surveying membrane landscapes: a new look at the bacterial cell surface. Nature Reviews Microbiology 21, 502–518 (2023).

23 Kumar, S. et al. Immobile lipopolysaccharides and outer membrane proteins differentially segregate in growing Escherichia coli. Proceedings of the National Academy of Sciences 122, e2414725122 (2025).

24 Rassam, P. et al. Supramolecular assemblies underpin turnover of outer membrane proteins in bacteria. Nature 523, 333–336 (2015).

25 Bergmiller, T. et al. Biased partitioning of the multidrug efflux pump AcrAB-TolC underlies long-lived phenotypic heterogeneity. Science 356, 311–315 (2017).

26 Gunasinghe, S. D. et al. The WD40 protein BamB mediates coupling of BAM complexes into assembly precincts in the bacterial outer membrane. Cell reports 23, 2782–2794 (2018).

27 Mühlradt, P. F., Menzel, J., Golecki, J. R. & Speth, V. Lateral mobility and surface density of lipopolysaccharide in the outer membrane of Salmonella typhimurium. European journal of biochemistry 43, 533–539 (1974).

28 Schindler, M., Osborn, M. & Koppel, D. E. Lateral diffusion of lipopolysaccharide in the outer membrane of Salmonella typhimurium. Nature 285, 261–263 (1980).

29 de Pedro, M. A., Grünfelder, C. G. & Schwarz, H. Restricted mobility of cell surface proteins in the polar regions of Escherichia coli. Journal of bacteriology 186, 2594–2602 (2004).

30 Nabarro, J. et al. Lipopolysaccharide lateral mobility in the Gram-negative bacterial outer membrane is confined and governed by interactions within the conserved Lipid A anchor. bioRxiv, 2025.2003. 2010.642448 (2025).

31 Verhoeven, G. S., Dogterom, M. & den Blaauwen, T. Absence of long-range diffusion of OmpA in E. coli is not caused by its peptidoglycan binding domain. BMC microbiology 13, 66 (2013).

32 Singer, S. J. & Nicolson, G. L. The Fluid Mosaic Model of the Structure of Cell Membranes: Cell membranes are viewed as two-dimensional solutions of oriented globular proteins and lipids. Science 175, 720–731 (1972).

33 Budin, I. et al. Viscous control of cellular respiration by membrane lipid composition. Science 362, 1186–1189 (2018).

34 Nikaido, H. Molecular basis of bacterial outer membrane permeability revisited. Microbiology and molecular biology reviews 67, 593–656 (2003).

35 Bengoechea, J.-A., Brandenburg, K., Seydel, U., Díaz, R. & Moriyón, I. Yersinia pseudotuberculosis and Yersinia pestis show increased outer membrane permeability to hydrophobic agents which correlates with lipopolysaccharide acyl-chain fluidity. Microbiology 144, 1517–1526 (1998).

36 Ginez, L. D. et al. Changes in fluidity of the E. Coli outer membrane in response to temperature, divalent cations and polymyxin-B show two different mechanisms of membrane fluidity adaptation. The FEBS Journal 289, 3550–3567 (2022).

37 Dumont, A., Malleron, A., Awwad, M., Dukan, S. & Vauzeilles, B. Click-mediated labeling of bacterial membranes through metabolic modification of the lipopolysaccharide inner core. Angewandte Chemie International Edition 13, 3143–3146 (2012).

38 Malinverni, J. C. & Silhavy, T. J. An ABC transport system that maintains lipid asymmetry in the gram-negative outer membrane. Proceedings of the National Academy of Sciences 106, 8009–8014 (2009).

39 Wotherspoon, P. et al. Structure of the MlaC-MlaD complex reveals molecular basis of periplasmic phospholipid transport. Nature Communications 15, 6394 (2024).

40 Kirschbaum, C. et al. Following phospholipid transfer through the OmpF3– MlaA–MlaC lipid shuttle with native mass spectrometry. Proceedings of the National Academy of Sciences 122, e2420041122 (2025).

41 Chong, Z. S., Woo, W. F. & Chng, S. S. Osmoporin OmpC forms a complex with MlaA to maintain outer membrane lipid asymmetry in E scherichia coli. Molecular microbiology 98, 1133–1146 (2015).

42 Thong, S. et al. Defining key roles for auxiliary proteins in an ABC transporter that maintains bacterial outer membrane lipid asymmetry. Elife 5, e19042 (2016).

43 Yeow, J. et al. The architecture of the OmpC–MlaA complex sheds light on the maintenance of outer membrane lipid asymmetry in Escherichia coli. Journal of Biological Chemistry 293, 11325–11340 (2018).

44 Ruiz, N., Falcone, B., Kahne, D. & Silhavy, T. J. Chemical conditionality: A genetic strategy to probe organelle assembly. Cell 121, 307–317 (2005).

45 Wu, T. et al. Identification of a multicomponent complex required for outer membrane biogenesis in Escherichia coli. Cell 121, 235–245 (2005).

46 Sutterlin, H. A. et al. Disruption of lipid homeostasis in the Gram-negative cell envelope activates a novel cell death pathway. Proceedings of the National Academy of Sciences 113, E1565–E1574 (2016).

47 Raetz, C. R. & Whitfield, C. Lipopolysaccharide endotoxins. Annual review of biochemistry 71, 635–700 (2002).

48 Fitzmaurice, D. R., Amador, A., Starr, T., Hocky, G. M. & Rojas, E. R. β-Barrel proteins dictate the effect of core oligosaccharide composition on outer membrane mechanics. Biophysical journal 124, 765–777 (2025).

49 Heinrichs, D. E., Yethon, J. A. & Whitfield, C. Molecular basis for structural diversity in the core regions of the lipopolysaccharides of Escherichia coli and Salmonella enterica. Molecular microbiology 30, 221–232 (1998).

50 Storek, K. M. et al. Monoclonal antibody targeting the β-barrel assembly machine of Escherichia coli is bactericidal. Proceedings of the National Academy of Sciences 115, 3692–3697 (2018).

51 Baba, T. et al. Construction of Escherichia coli K-12 in-frame, single-gene knockout mutants: the Keio collection. Molecular systems biology 2, 2006.0008 (2006).

52 Bentley, A. T. & Klebba, P. E. Effect of lipopolysaccharide structure on reactivity of antiporin monoclonal antibodies with the bacterial cell surface. Journal of bacteriology 170, 1063–1068 (1988).

53 Coleman Jr, W. G. & Leive, L. Two mutations which affect the barrier function of the Escherichia coli K-12 outer membrane. Journal of bacteriology 139, 899–910 (1979).

54 Kneidinger, B. et al. Biosynthesis pathway of ADP-L-glycero-β-D-manno-heptose in Escherichia coli. Journal of Bacteriology 184, 363–369 (2002).

55 Yethon, J. A. & Whitfield, C. Purification and characterization of WaaP from Escherichia coli, a lipopolysaccharide kinase essential for outer membrane stability. Journal of Biological Chemistry 276, 5498–5504 (2001).

56 Konovalova, A., Kahne, D. E. & Silhavy, T. J. Outer membrane biogenesis. Annual review of microbiology 71, 539–556 (2017).

57 Dekker, N. Outer-membrane phospholipase A: known structure, unknown biological function: MicroReview. Molecular microbiology 35, 711–717 (2000).

58 Bishop, R. E. Structural biology of membrane-intrinsic β-barrel enzymes: Sentinels of the bacterial outer membrane. Biochimica et Biophysica Acta (BBA)-Biomembranes 1778, 1881–1896 (2008).

59 Kramer, R., Zandwijken, D., Egmond, M. R. & Dekker, N. In vitro folding, purification and characterization of Escherichia coli outer membrane protease OmpT. European journal of biochemistry 267, 885–893 (2000).

60 Li, G.-W., Burkhardt, D., Gross, C. & Weissman, J. S. Quantifying absolute protein synthesis rates reveals principles underlying allocation of cellular resources. Cell 157, 624–635 (2014).

61 Ekholm, F. et al. Synthesis of the copper chelator TGTA and evaluation of its ability to protect biomolecules from copper induced degradation during copper catalyzed azide–alkyne bioconjugation reactions. Organic & Biomolecular Chemistry 14, 849–852 (2016).

62 Ursell, T. et al. Rapid, precise quantification of bacterial cellular dimensions across a genomic-scale knockout library. BMC biology 15, 1–15 (2017).

63 Sliusarenko, O., Heinritz, J., Emonet, T. & Jacobs-Wagner, C. High-throughput, subpixel precision analysis of bacterial morphogenesis and intracellular spatio-temporal dynamics. Molecular microbiology 80, 612–627 (2011).

64 Van Valen, D. A. et al. Deep learning automates the quantitative analysis of individual cells in live-cell imaging experiments. PLoS computational biology 12, e1005177 (2016).

65 Neidhardt, F. C., Bloch, P. L. & Smith, D. F. Culture medium for enterobacteria. Journal of bacteriology 119, 736–747 (1974).

66 Li, X.-t., et al. tCRISPRi: tunable and reversible, one-step control of gene expression. Scientific reports 6, 39076 (2016).

67 Silvis, M. R. et al. Morphological and transcriptional responses to CRISPRi knockdown of essential genes in Escherichia coli. MBio 12, 10.1128/mbio.02561-02521 (2021).

68 Jo, S., Kim, T., Iyer, V. G. & Im, W. CHARMM-GUI: a web-based graphical user interface for CHARMM. Journal of computational chemistry 29, 1859–1865 (2008).

69 Li, Y., Liu, J. & Gumbart, J. C. in Structure and Function of Membrane Proteins 237–251 (Springer, 2021).

70 Wu, E. L. et al. E. coli outer membrane and interactions with OmpLA. Biophysical journal 106, 2493–2502 (2014).

71 Balusek, C. & Gumbart, J. C. Role of the native outer-membrane environment on the transporter BtuB. Biophysical Journal 111, 1409–1417 (2016).

72 Klauda, J. B. et al. Update of the CHARMM all-atom additive force field for lipids: validation on six lipid types. The journal of physical chemistry B 114, 7830–7843 (2010).

73 Phillips, J. C. et al. Scalable molecular dynamics on CPU and GPU architectures with NAMD. The Journal of chemical physics 153 (2020).

74 Balusek, C. et al. Accelerating membrane simulations with hydrogen mass repartitioning. Journal of chemical theory and computation 15, 4673–4686 (2019).

75 Humphrey, W., Dalke, A. & Schulten, K. VMD: visual molecular dynamics. Journal of molecular graphics 14, 33–38 (1996).

76 Virtanen, P. et al. SciPy 1.0: fundamental algorithms for scientific computing in Python. Nature methods 17, 261–272 (2020).

77 Leive, L., Shovlin, V. K. & Mergenhagen, S. E. Physical, chemical, and immunological properties of lipopolysaccharide released from Escherichia coli by ethylenediaminetetraacetate. J Biol Chem 243, 6384–6391 (1968).

78 Button, J. E., Silhavy, T. J. & Ruiz, N. A suppressor of cell death caused by the loss of σE downregulates extracytoplasmic stress responses and outer membrane vesicle production in Escherichia coli. Journal of bacteriology 189, 1523–1530 (2007).

